# The antioxidant networks act as a firewall in maintenance of genomic integrity

**DOI:** 10.1101/2025.08.11.669663

**Authors:** Hongen Geng, Xiaoping Huang, Yiyu Cheng, Hanqi Chang, Yi Wang, Yuanzhuo Yu, Yunfeng Zhang, Liuxing Yang, Chunrong Liu, Yao Zhao, FuYi Wang, Xiang Li, Changlin Liu

## Abstract

Reactive oxygen species (ROS) threatens genomic integrity by both destruction of DNA metabolic enzymes and oxidative DNA damage. We find a negative correlation between oxidative DNA damage and activity of antioxidant enzymes. An antioxidant network formed by interactions between the antioxidant enzymes and the proteins involved in maintenance of genomic integrity was indentified in the nucleus by multi-proteomic investigations. The protein-protein interactions in the antioxidant network are up-regulated under oxidative stress, as proved by quantitative proximity ligation assays *in situ* within single cells. The antioxidant network catalyzes the ROS dismutation cascade before ROS attacks DNA-processing enzymes, thereby preventing DNA and DNA metabolic enzymes from oxidative damage and dysfunction. Therefore, the antioxidant network plays a firewall role in maintenance of genomic integrity.

## Introduction

The accumulation of reactive oxygen species (ROS, here designated mainly as hydrogen peroxide, H_2_O_2_; superoxide anion, O_2_^•-^; and hydroxyl radical, •OH) within a cell nucleus is stemmed not only from its mitochondria (1,2), but also from its nuclear self-generation by oxidative demethylation of both methylated histones and methylated DNA (3,4), and by catalytic processes of enzymes such as ribonucleotide reductase (5), NADPH oxidase 4 (6) and spermine oxidase (7). The nuclear ROS, regardless of its sources, can lead to highly frequent destruction of genomic integrity (8–15) that is linked to many human diseases, e.g., cancer, neurodegenerative disorders and aging (16–21). Of particular notes is that the demethylation of histones and CpGs in gene promoters and other regulatory regions generates ROS that oxidize DNA (11). Hydroxyl radical (•OH) is responsible for oxidative DNA damage (ODD) (10), and 8-oxo-7,8-dihydro-2^’^- deoxyguanosine (8-oxodG) is its major oxidation product, because guanine among the DNA nucleobases is most susceptible to attack by ROS (22–26). Only the Fenton reaction between H_2_O_2_ and redox-active Fe^2+^ can generate the most reactive •OH (10). The redox-active Fe-S cluster [4Fe-4S]^2+^ plays a critical role in DNA metabolism (27). The possible reaction of H_2_O_2_ with [4Fe-4S]^2+^ in the nucleus can not only generate •OH, but also result in dysfunction of the [4Fe-4S]^2+^- containing DNA metabolic enzymes essential for maintenance of nuclear genomic integrity (28). This implies that the [4Fe-4S]^2+^ in the DNA metabolic enzymes plays an essential role in ODD, and ODD occurs in an extremely high frequency in the nucleus. Moreover, ROS can also cause DNA replication stress and even collapse of the DNA replication fork (29,30). However, the fact is that the frequency of endogenous ODD estimated by 8-oxodG only approaches 4% of the total damage frequency (∼ 70000 lesions) to which the DNA in each human cell is subject per day (31,32). This raises a question of whether cells had acquired a defense mechanism during evolution that eliminates the excessive ROS that threatens genomic integrity in the nucleus. We reasoned that the antioxidant enzymes, including Cu/Zn superoxide dismutase (SOD1), catalase (CAT), and peroxiredoxin family (PRDXs), could play a crucial role in this defense mechanism unidentified so far, because SOD1 catalyzes the conversion of O_2_^•-^ into H_2_O_2_, whereas CAT and PRDXs promote the decomposition of H_2_O_2_. These antioxidant enzymes were displayed higher distributions in the nucleus as ROS levels rise in the cell (33–37). The single antioxidant enzymes including SOD1, PRDX1/2 and CAT were respectively found to involve in maintenance of genomic integrity in the cell nuclei of different species (5,33,34,39–47). Thus, we propose that these antioxidant enzymes could form an antioxidant network with the proteins essential for maintaining genomic integrity in the nucleus, functioning to eliminate excessive ROS through catalyzing the dismutation cascade O_2_^•-^→H_2_O_2_→H_2_O in the defense mechanism.

To test our proposal, the antioxidant enzymes SOD1, CAT, and PRDX2 were selected because of their involvement in maintenance of genomic integrity (5,33,34,39–47). First, we found that the antioxidant enzymes not only reduce the oxidative DNA cleavage caused by the mixtures of [4Fe-4S]^2+^-containing primase and H_2_O_2_ in solution, but also show a negative correlation between their activity and cellular ODD. Then, a huge amount of multi-proteomic data from both the whole cell and the nucleus, produced by affinity purification- and co-immunoprecipitation-coupled quantitative mass spectrometry (AP- and CoIP-qMS), were integrated, together with single-cell *in situ* proximity ligation assays (PLAs) and Western blotting tests, under H_2_O_2_-untreated and - treated conditions. The results reveal the presence of interaction networks not only among the antioxidant enzymes but also between the antioxidant enzymes and the enzymes essential for maintenance of genomic integrity in both whole cells and the nucleus. The networks assembled by the protein-protein interactions (PPIs) are referred respectively to as the antioxidant network in the cell and the antioxidant subnetwork in the nucleus. The PPIs, not abundance of the proteins, in the antioxidant subnetworks in the nucleus are notably up-regulated upon H_2_O_2_ treatment of the cell. The antioxidant subnetworks can act as a firewall that safeguards nuclear DNA, and might be an alternative mechanism for the cell to maintain its nuclear genomic integrity in addition to DNA damage response (48–50).

## Results

### Oxidative DNA cleavage is reduced by the antioxidant enzymes in solution

To examine the possibility that the reaction of H_2_O_2_ with [4Fe-4S]^2+^ triggers oxidative DNA cleavage, the human primase subunit PRIM2 that only contains the iron cofactor [4Fe-4S]^2+^ was expressed, reconstructed and characterized. ICP(inductively coupled plasma)-MS determinations indicated the presence of iron and UV-visible absorption spectra confirmed that the iron exists in the form of [4Fe-4S]^2+^ in the enzyme (51) (fig. S1A). Both the reaction of H_2_O_2_ with [4Fe-4S]^2+^ and the exposure of [4Fe-4S]^2+^ to air all can generate •OH that triggers oxidative DNA cleavage and lead to destruction or oxidation of the cluster, because O_2_ in air can react with the cluster (52–54). To remove the DNA cleavage portion mediated by the reaction of [4Fe-4S]^2+^ with O_2_ from the total DNA cleavage, the oxidative cleavage of the supercoiled plasmid DNA (pRSETA) was observed in the glovebox filled with argon gas. The results showed that the proportions of the supercoiled (**I**) and nicked/linear DNA (**Ⅱ+Ⅲ**) were reduced and increased respectively with increasing PRIM2 and H_2_O_2_ concentrations (Fig. 1A). Moreover, the amount of three forms of the DNA was changed with incubation time at the given concentrations of PRIM2 and H_2_O_2_ (fig. S1B). In comparison, oxidative DNA cleavage was examined with ferrous sulfate whose concentration was equal to four times of the PRIM2 concentration and to the iron concentration in PRIM2 under the same conditions. The result indicated that the oxidative DNA cleavage in the same incubation periods was much more in the Fe(II)-containing solution than in the PRIM2-containing solution (fig. S1C). Thus, the DNA cleavage caused by PRIM2 was much less in efficiency than that by the Fe(II) salt, likely it is difficult for H_2_O_2_ to access to the [4Fe-4S]^2+^ that is deeply buried inside the protein. Evidently, these results support the hypothesis that the reaction of H_2_O_2_ with [4Fe-4S]^2+^ in the DNA metabolic enzymes results in oxidative DNA cleavage.

**Fig. 1.**
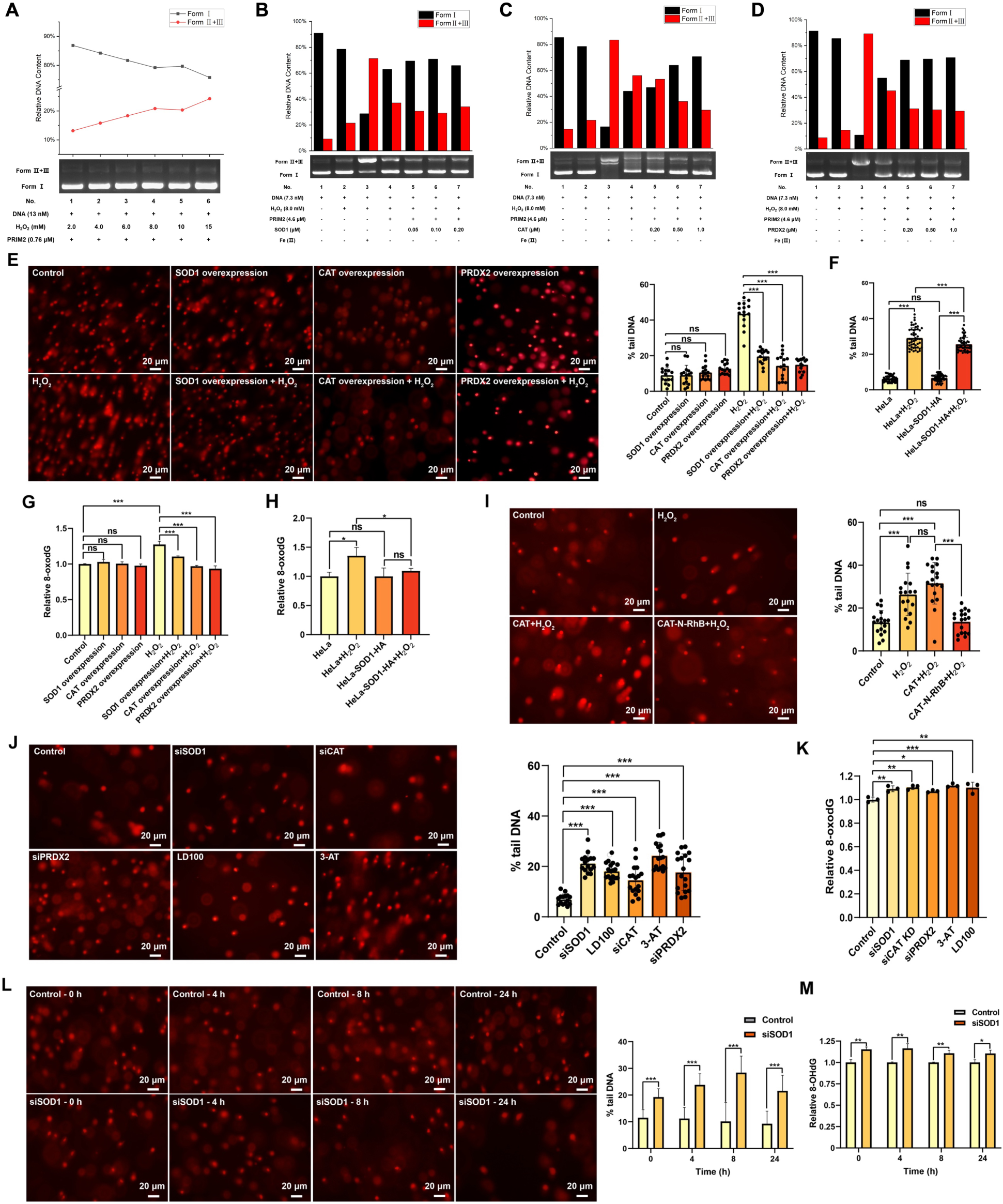
Oxidative DNA damage is modulated by antioxidant enzymes *in vitro* and *in vivo*. **A**) Agarose gel electrophoresis showed that H_2_O_2_ (2-15 mM) mediates concentration-dependent cleavage of plasmid DNA, in the presence of PRIM2 protein containing [4Fe-4S]^2+^. **B-D**) Agarose gel electrophoresis showed that SOD1 (**B**, 0.05-0.20 μM), CAT (**C**, 0.20-1.00 μM), and PRDX2 (**D**, 0.20-1.00 μM) suppress H_2_O_2_-mediated plasmid DNA cleavage, in the presence of PRIM2 protein containing [4Fe-4S]^2+^. Representative electrophoresis images are presented, and quantitative results were derived from grayscale analysis of DNA bands (**B-D**). **E-H**) Comet tail and 8-oxodG assays showed that oxidative DNA damage is suppressed by elevating activity of the antioxidant enzymes. HeLa cells were transfected with plasmids (**E**, **G**) or infected by lentivirus (**F**, **H**), treated with or without 400 μM H_2_O_2_, and harvested for comet and 8-oxodG assays. **I**) Comet tail assays showed that oxidative DNA damage is suppressed by intracellular delivery of CAT with N-terminal rhodamine B modification (CAT-N-RhB). Here, A549 cells were used. **J**, **K**) Comet tail and 8-oxodG assays showed that reducing antioxidant enzyme activity elevates oxidative DNA damage. Following transfected with siRNAs or incubated with the SOD1 (50 μM **LD100**) or CAT (10 mM **3-AT**) inhibitor, HeLa cells were harvested for comet tail and 8-oxodG assays. **L**, **M**) The levels of both DNA comet tail and 8-oxodG were increased with knockdown time of SOD1. Following transfected with siRNAs of SOD1 for 0-24 h, HeLa cells were harvested for comet tail and 8-oxodG assays. Comet tail content was analyzed using Comet Score 2.0 (n ≥ 15); scale bar: 20 μm; n≥50 for comet assays; n=3 for 8-oxodG assays. \**P* < 0.05, \*\**P* < 0.01, \*\*\**P* < 0.001; One-way ANOVA. All error bars are SD.

We speculate that the presence of the antioxidant enzymes in the solution that promote H_2_O_2_ decomposition or bind to DNA could prevent DNA from the oxidative cleavage. Therefore, the conversion of the supercoiled DNA into its nicked and linear forms was examined respectively in the SOD-, CAT- and PRDX2-containing solutions under the air-isolated condition. Form **I** and **Ⅱ+Ⅲ** of the DNA were respectively 63% and 37% following incubation for 40 min with 4.6 μM PRIM2 and 8 mM H_2_O_2_, and were continuously elevated and reduced with increased addition of SOD1 (Fig. 1B). Furthermore, we found that the treatment with CAT and PRDX2 also resulted in significantly elevated amount of supercoiled DNA and reduced amount of nicked and linear DNA (Fig 1, C and D). Evidently, these results demonstrated that the presence of three antioxidant enzymes can significantly mitigate the oxidative DNA cleavage caused by the reaction between H_2_O_2_ and PRIM2. However, the antioxidant enzymes prevent DNA from the oxidative cleavage likely by different mechanisms. CAT catalyzes H_2_O_2_ decomposition at a high rate (55), and notably decrease the amount of H_2_O_2_ that reacts with [4Fe-4S]^2+^ in PRIM2. SOD1 prevents oxidative attacks from ROS only by binding to DNA (34,45) or likely by binding to the [4Fe-4S]^2+^ enzymes, thereby safeguarding both the DNA or the enzymes, as previously reported (56,57). On the other hand, PRDX2 not only accelerates H_2_O_2_ decomposition (36), but also binds to DNA in a decameric form (5).

Consequently, oxidative DNA cleavage can occur in the solution containing H_2_O_2_ and the [4Fe-4S]^2+^ enzymes, just as inferred generally (28). The antioxidant enzymes can significantly reduce oxidative DNA cleavage both by preventing the reaction of H_2_O_2_ with [4Fe-4S]^2+^ in the enzymes and by the association with DNA in the solution.

### Oxidative DNA damage is suppressed by the antioxidant enzymes in the cell

The reduction of oxidative DNA cleavage by the antioxidant enzymes in solution inspired us to examine whether the antioxidant enzymes can mitigate oxidative damage of nuclear genomes in the cell. The ODD change with addition of H_2_O_2_ was firstly examined within HeLa cells. The cell viability and the intracellular H_2_O_2_ content were observed to be reduced and increased respectively with both increased addition of H_2_O_2_ and elongation of incubation periods (fig. S2). The cellular content of comet tails and 8-oxodG, a pair of established biomarkers for ODD that were determined respectively by gel electrophoresis (comet assay) and ELISA assays (58), was found to rise with addition of and incubation with H_2_O_2_ (0-400 μM for 0-6 h, fig. S3). The cells still maintained more than 80% of viability but displayed significant ODD under the conditions treated for 4 h with 400 μM H_2_O_2_. Therefore, the following tests were performed all in the cells under this condition. Then, the nuclear distributions of the antioxidant enzymes SOD1, CAT and PRDX2 were confirmed in the cells (fig. S4), as reported previously (34–37, 59).

The ODD changes in the nucleus with activity of the antioxidant enzymes was examined under H_2_O_2_-untreated and -treated conditions. Overexpression and cell delivery (60) up-regulated intracellular content and activity of the antioxidant enzymes, whereas knockdown and addition of the specific inhibitor LD100 of SOD1 (61) or 3-AT (1,2,4-Triazole) of CAT (62) down-regulated their content and activity, as indicated respectively by increased and decreased intracellular H_2_O_2_ levels (fig. S5). The quantification of comet tails and 8-oxodG indicated that treatment for 4 h with H_2_O_2_ led respectively to 5- and 1.27-fold increase of comet tails (from 5% to > 40%) and 8-oxodG (from 100% to ∼ 130%) (Fig. 1, E to H). On the one hand, the overexpression of SOD1, CAT and PRDX2 each suppressed the H_2_O_2_-induced formation of both comet tails (from 43% to 14-19%) and 8-oxodG (from 127% to 93-110%) (Fig. 1, E to H). It is notable that elevating cellular SOD1 levels, whether overexpressed by lentivirus infection or Lipofactamine transfection, all inhibited ODD, although SOD1 increases cellular H_2_O_2_ content by catalyzing dismutation of O_2_^•-^ into H_2_O_2_ (Fig. 1, F and H). However, the enzymes that reduce cellular H_2_O_2_ content more significantly suppressed ODD than SOD1 (Fig. 1, E and G). Furthermore, intracellular delivery of active CAT (60) also inhibited the formation of comet tails (from 31% to 14%) (Fig. 1I). These results indicated that the ODD mediated by addition of H_2_O_2_ was considerably reduced with elevating activity of the antioxidant enzymes. On the other hand, the respective siRNA interference of the genes *SOD1*, *CAT* and *PRDX2* resulted in elevated comet tail content from 7.0% to 14-21% and 8-oxodG levels from 100% to 110% (Fig. 1, J and K). The addition of H_2_O_2_ further increased the content of the ODD markers in the SOD1 knockdown cells, as expected (Fig. 1, L and M). Moreover, comet tails and 8-oxodG of DNA were observed to increase with knockdown time of SOD1 in cells (Fig. 1, L and M). The respective inhibition of SOD1 and CAT also notably raised comet tails from 7.0% to 18-24%, and 8-oxodG content from 100% to 110% (Fig. 1, J and K). Obviously, this elevated DNA damage stems from the increased cellular ROS levels caused by decreasing activity of the antioxidant enzymes.

In brief, ODD in the nucleus was suppressed by elevating activity of the three antioxidant enzymes *via* their overexpression or carrier-free cell delivery, but was elevated by reducing their activity through their knockdown or specific inhibition. These results indicate that the antioxidant enzymes are not only a guardian of nuclear genomic integrity, but also might protect the [4Fe-4S]^2+^ in the proteins responsible for maintenance of genomic integrity, which is in agreement with the previously reported compartment-specific protection of Fe-S proteins by superoxide dismutase (56).

### Reducing SOD1 activity impacts the function of the proteins essential for maintenance of genomic integrity

To support the conclusion that elevating activity of the antioxidant enzymes is essential for suppression of ODD in the nucleus, the whole-cell proteomic data were collected and compared by data independent acquisition (DIA)-qMS for the H_2_O_2_-untreated and -treated groups of HeLa cells with undisturbed and reduced SOD1 expression (Fig. 2A). DIA quantifies proteins with comparable precision as data dependent acquisition (DDA), while exhibiting broader protein coverage and enhanced data reproducibility (63–65). A total of 5040 proteins were quantified in the 4 groups of cells, and of them, 4353 proteins (86% of the total) were shared by 12 tests (Fig. 2, A and B), and the unique proteins were less than 100 for each group of cells, demonstrating the high reproducibility of the DIA-based quantitative proteomic data. Principal component analysis (PCA) showed that the MS data closely approached within each group, but were well separated between the groups (Fig. 2C). These results indicated that the differentials in protein expression were significant among the groups and the reproducibility was high for biological replicates. Meanwhile, a total of 4818 proteins with high confidence were identified. Of them, 2400 proteins displayed significant differentials in expression at least between two sets of test conditions and were well clustered inside each group (fig. S6A). The clustering data showed that the proteomic data are high reproducible inside each group and notably differential between two groups, as indicated by PCA. Therefore, the regions determined by PC1 and PC2 suggested that SOD1 knockdown more prominently impacts protein expression than treatment with H_2_O_2_ inside the cells (Fig. 2C).

**Fig. 2.**
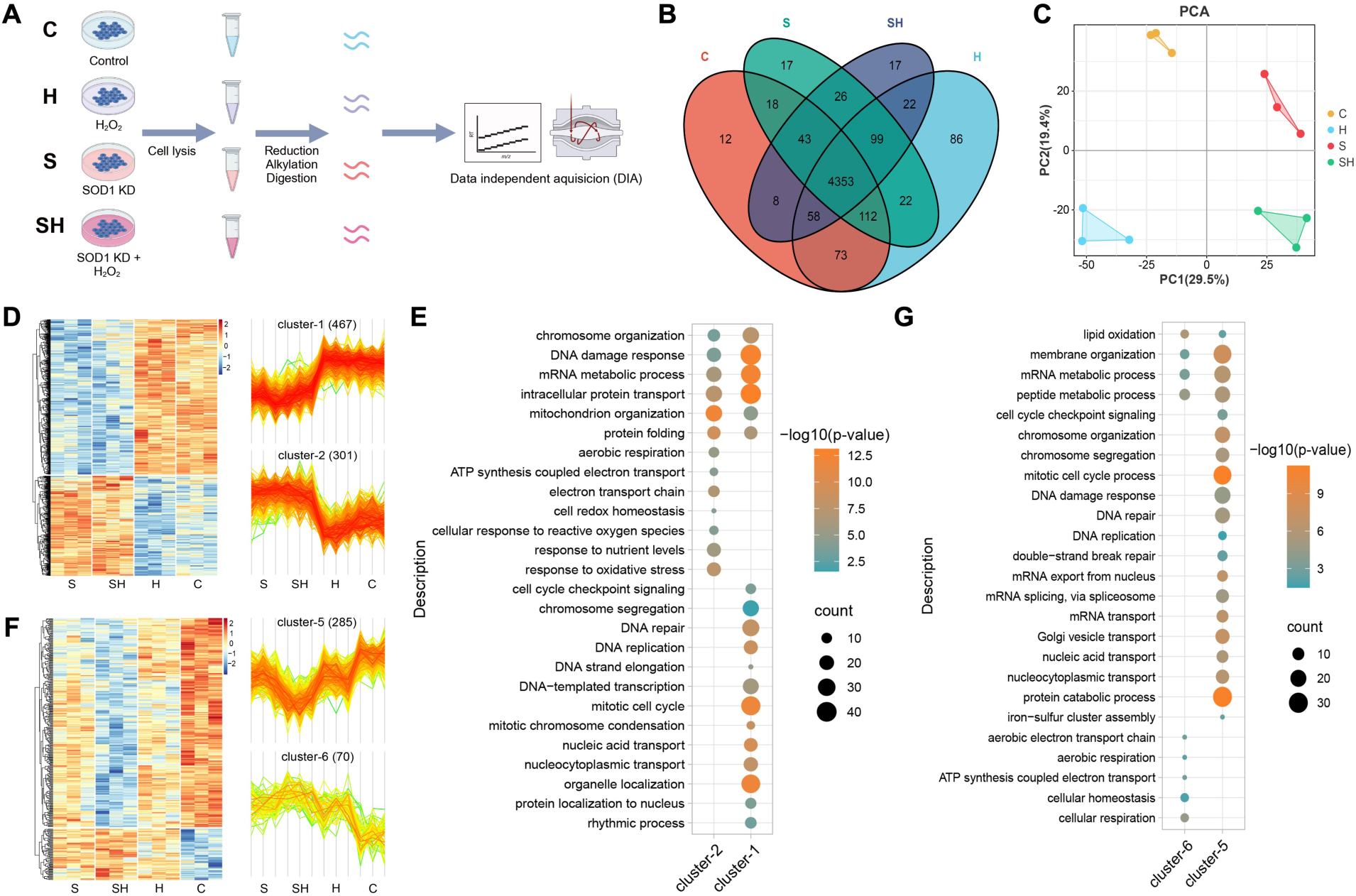
SOD1 impacts the function of key proteins involved in maintaining genomic integrity. **A**) Flow diagram of the integrative proteomic analysis evaluating the effects of SOD1 knockdown and H_2_O_2_ stimulation on cellular processes (**C**: control, **H**: H_2_O_2_ treatment, **S**: SOD1 knockdown, **SH**: SOD1 knockdown and H_2_O_2_ treatment). Proteomic data was acquired using a Data-Independent Acquisition (DIA) method. **B**) Venn diagram showing the overlap of proteins with quantitative information. **C**) PCA analysis of protein expression profiles. **D**, **E**) Clustering (**D**) and GO analysis (**E**) of proteins specifically responsive to SOD1 knockdown. **F**, **G**) Clustering (**F**) and GO analysis (**G**) of proteins responsive to both SOD1 knockdown and H_2_O_2_ stimulation. Quantitative proteomics was performed in triplicate. Protein abundances were normalized based on MS signal intensity (**D**, **F**).

The heat map obtained by clustering showed that the differentials in their quantitative proteomic information are noteworthy between these clusters (fig. S6A). The proteins responded to the reduced SOD1 activity were first selected and clustered, whose expression was respectively down-regulated (467, cluster-1) and up-regulated (301, cluster-2) in the SOD1 knockdown cells (Fig. 2D). These proteins were found to display significant differentials only between SOD1 expression-undisturbed (C and H groups) and -reduced (S and SH groups) cells, revealing that the abundance of the proteins is impacted by SOD1 knockdown, but not by H_2_O_2_ treatment. The proteins in cluster-1 are involved in nuclear functions, including DNA replication, DNA damage response and repair, transport and metabolism of nucleic acids, condensation, organization and segregation of chromosomes, cell cycle, and protein localization to the nucleus (Fig. 2E). Obviously, these nuclear functions essential for maintenance of genomic integrity are impaired by the reduced SOD1 activity. On the other hand, the proteins in cluster-2 mainly are involved in the mitochondrial functions, including aerobic respiration, electron transfer and responses to oxidative stress and ROS, and mitochondrion organization (Fig. 2E), indicating that the reduced SOD1 activity disturbs mitochondrial ROS metabolism. The proteins sensitive to H_2_O_2_ treatment were found to have differential expression in H_2_O_2_-treated and control groups, and clustered into cluster-3 (79 proteins, down-regulated) and -4 (114 proteins, up-regulated) (fig. S6B). The proteins (193) in response to H_2_O_2_ treatment were much less than those (768) in response to SOD1 knockdown. Moreover, these proteins do not participate in the nuclear functions other than cluster-3 that involves only in regulation of cell cycle (fig. S6C). These results allowed us to speculate that the elevated H_2_O_2_ level does not exert significant impact on the functions of many proteins essential for maintenance of genomic integrity.

To prove this speculation, the proteins, which were up-regulated and down-regulated by dual interference of SOD1 knockdown and H_2_O_2_ treatment, were respectively clustered into cluster-5 and -6 (Fig. 2F). Although their expression displayed significant differentials compared to the control group, the number of the proteins covered by cluster-5 and -6 were much less than that by cluster-1 and -2 (Fig. 2D). As indicated by GO enrichment, the down-regulated cluster-5 (285 proteins) involved in the same nuclear functions essential for maintenance of genomic integrity as cluster-1, the up-regulated cluster-6 (70 proteins) was independent of the nuclear functions (Fig. 2G). These results indicate that only reducing SOD1 activity in the dual interference has a significant impact on the essential functions in the nucleus. In summary, reducing intracellular SOD1 activity compromises the functions of the nuclear proteins essential for maintenance of genomic integrity and disturbs mitochondrial ROS metabolism. Increasing intracellular H_2_O_2_ levels brings about the effect similar to reducing SOD1 activity, but exerts much weaker impacts on the functions of the maintenance proteins of genomic integrity than reducing SOD1 activity.

### The interactions between the antioxidant enzymes and nuclear proteins are up-regulated by H_2_O_2_ treatment

The activity of the antioxidant enzymes is negatively correlated with ODD, and plays a crucial role in maintenance of nuclear genomic integrity, suggesting that the antioxidant enzymes interact extensively with the maintenance proteins of genomic integrity. Thus, we quantitatively identified the antioxidant enzymes-involved PPIs by the DIA-qMS scheme coupled with CoIP in the H_2_O_2_-untreated and -treated cells (Fig. 3A).

**Fig. 3.**
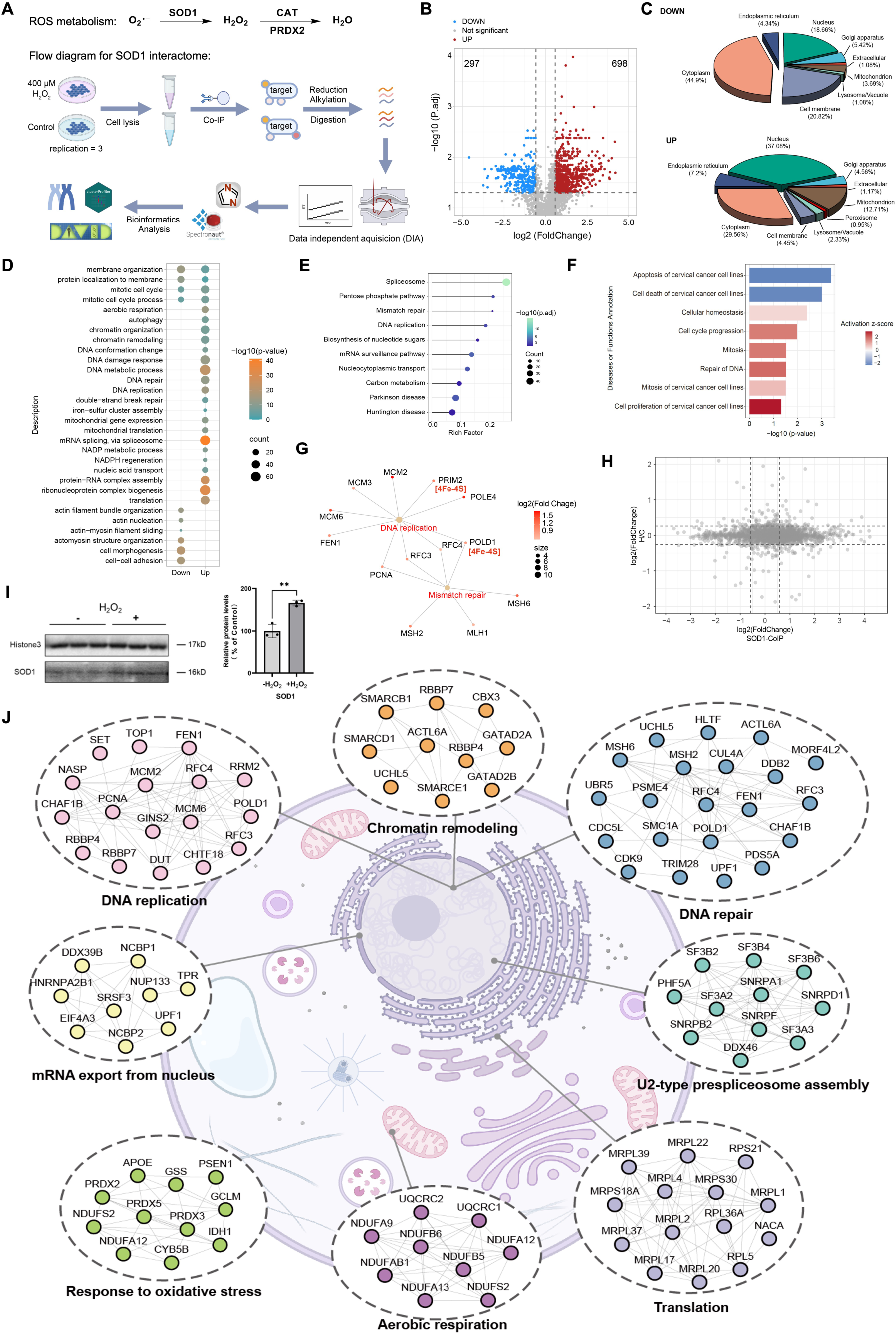
H_2_O_2_ enhances interactions between antioxidant enzymes and nuclear proteins. **A**) Flow diagram of SOD1 interactome by CoIP coupled DIA-qMS. ROS metabolism involving SOD1, CAT and PRDX2 is also shown. **B**) Volcano plot showing the proteins whose interaction with SOD1 are significantly altered by H_2_O_2_ treatment (p. adjust ≤ 0.05, fold change ≥ 1.5). **C**, **D**, **F**) Subcellular localization (**C**), GO analysis (**D**), and IPA annotation (**F**) of the proteins whose PPIs with SOD1 are up- and down-regulated by H_2_O_2_ treatment. **E**) KEGG pathway analysis of the proteins involved in the enhanced interactions with SOD1 by H_2_O_2_ treatment. **G**) Proteins involved in DNA replication and mismatch repair pathways that display enhanced interactions with SOD1 after H_2_O_2_ treatment. **H**) Integrative analysis of the global protein quantification (y-axis) and the SOD1 interactome data (x-axis). **I**) Western blotting demonstrated that oxidative stress enhances the binding of SOD1 to chromatin in the nucleus. **J**) The landscape mapping of the up-regulated PPIs with SOD1 by H_2_O_2_ treatment.

The interactomes of SOD1, CAT and PRDX2 were respectively identified by CoIP-DIA-qMS (Fig. 3A). The PCA distributions of each enzyme-involved PPI data indicated that these quantitative proteomic data displayed significant differences between the H_2_O_2_-untreated and - treated samples (fig. S7A). The pairwise comparisons showed that these proteomic data were well reproduced respectively under the two kinds of tested conditions (fig. S7B).

First, a total of 2773 proteins were identified to interact with SOD1 in H_2_O_2_-untreated and - treated cells, 995 proteins of which display significant differences in PPIs (p. adjust ≤ 0.05, fold change ≥ 1.5) (Fig. 3B). The database BioGRID reported 409 SOD1-interacting proteins based on the reported results (66), 253 (63%) proteins of which were identified in this study, indicating that the unbiased deep identification based on our workflow can cover more SOD1-interacting proteins than those reported previously. Of 995 proteins, the interactions of 693 and 297 proteins with SOD1 were respectively up- and down-regulated upon H_2_O_2_ treatment (Fig. 3B). The subcellular localization indicated that the up-regulated PPIs occur mainly in the nucleus (Fig. 3C), which could be attributed to the H_2_O_2_-drived translocation of SOD1 to the nucleus (33,34), whereas the down-regulated PPIs appear mainly in the cytoplasm.

Functional and pathway annotation was conducted for the proteins whose interactions with SOD1 were altered upon H_2_O_2_ treatment. The proteins with the up-regulated PPIs were especially worthy of our attention, as indicated by GO enrichment. They play a crucial role in nuclear functions essential for maintenance of genomic integrity, e.g., DNA damage response, DNA replication, chromatin organization and remodeling, and cell cycle, whereas the proteins with the down-regulated PPIs are involved in limited functions, e.g., membrane organization (Fig. 3D). KEGG enrichment indicated that the proteins are key participants in the nuclear pathways including spliceosome, pentose phosphate pathway, DNA replication and repair (Fig. 3E). Furthermore, ingenuity pathway analysis (IPA) showed that the SOD1-involved PPIs elevated upon H_2_O_2_ treatment activate the cell survival and proliferation pathways (Fig. 3F, red bars) including cell cycle progression, mitosis, homeostasis, and DNA repair, but repress cell death and apoptosis (Fig. 3F, blue bars). Further analysis revealed that the SOD1 interactions with numerous enzymes required for DNA replication and repair, for example, PRIM2 (a DNA primase subunit) and POLD1 (a DNA polymerase subunit), were notably elevated upon H_2_O_2_ treatment (Fig. 3G). This pair of enzyme subunits essentially require the cluster [4Fe-4S]^2+^ for the functions of their active complexes (28, 67–69) and for maintenance of genomic stability (70).

The alterations in the SOD1-involved PPIs could be attributed to the changes in global abundance of SOD1-interacting proteins or to the intracellular H_2_O_2_-drived SOD1 translocation toward the nucleus, or to both. To eliminate the former possibility, the cell-wide proteins were relatively quantified by DIA-qMS (fig. S7C). The integrated analysis of this quantification, together with the SOD1 interactome CoIP-DIA-qMS data, indicated that 98% of the proteins whose interactions with SOD1 were altered upon H_2_O_2_ treatment did not display a significant alteration in abundance (Fig. 3H, fold change ≥ 1.5), which is well consistent with the conclusion that is stemmed from whole-cell proteomic tests under the conditions of undisturbed and reduced SOD1 expression (Fig. 2D). On the other hand, the amount of chromatin-bound SOD1 was elevated to 165% of the control following H_2_O_2_ treatment (Fig. 3I), indicating the translocation of numerous cytoplasmic SOD1 toward the nucleus upon oxidative stress. These results revealed that the H_2_O_2_-drived SOD1 translocation from the cytoplasm to the nucleus up-regulates the PPIs with SOD1 in the nucleus, but the cell-wide abundance of the SOD1-interected proteins is not altered upon H_2_O_2_ stimulation. In fact, the global organelle profiling disclosed that many proteins are regulated by changes in their spatial distribution rather than by changes in their total abundance (71).

The landscape mapping and clustering of the SOD1-involved PPIs revealed that the up-regulated PPIs upon H_2_O_2_ stimulation can assemble into a big network, which consists of nuclear and cytoplasmic modules (10 in total, Fig. 3J). The modules in the nucleus occupy half of the network and the two largest nuclear modules are responsible for DNA replication and repair. Collectively, these results indicated that the SOD1-involved up-regulated PPI modules, also including chromatin remodeling, are responsible for maintenance of nuclear genomic integrity.

Then, the PPIs with CAT and PRDX2 were respectively quantified under H_2_O_2_-untreated and - treated conditions. A total of 1675 proteins were found to have interactions with CAT, 278 proteins of which display significant differences in PPIs (p. adjust ≤ 0.05, fold change ≥ 1.5) (fig. S8A). H_2_O_2_ treatment resulted in much more up-regulated PPIs (261 proteins) than down-regulated PPIs (17 proteins). Although the PPIs with CAT were decreased (from 34% to 31%) in the nucleus (fig. S8B), CAT was still found to have up-regulated interactions with PRIM2, POLD1 and MCM2, as found for SOD1. The increased PPIs with CAT in the mitochondrion indicated an essential role of CAT in the counteraction of H_2_O_2_ stimulation. GO enrichment showed that the up-regulation in the PPIs is linked to the functions including mRNA splicing and export, DNA repair, cell cycle and division, and chromosome organization (fig. S8C). On the other hand, the interactions of 2529 proteins with PRDX2 were quantified, 452 proteins of which displayed significantly differential PPIs (p. adjust ≤ 0.05, fold change ≥ 1.5) (fig. S8D). H_2_O_2_ treatment elevated the interactions of 262 proteins with PRDX2, and reduced those of 190 proteins with PRDX2. Moreover, the PPIs with PRDX2 were up-regulated from 25.8% to 42.7% in the nucleus, and down-regulated from 22.2% to 2.5% in the mitochondrion, indicating that PRDX2 plays an essential role in the nucleus under oxidative stress (fig. S8E). In fact, GO analysis revealed that the PPIs with PRDX2 participate in execution of the functions including mRNA splicing, DNA repair and replication, cell cycle and DNA damage response (fig. S8F).

It is noteworthy that the elevated interactions also occur among the three antioxidant enzymes. Moreover, SOD1 was observed to have the elevated interaction with glyceraldehyde-3-phosphate dehydrogenase (GAPDH), likely because GAPDH is sensitive to reversible oxidative inactivation by H_2_O_2_, and plays a role in maintenance of DNA integrity and oxidative stress response (72).

### The antioxidant enzymes share the subnetworks consisted of the up-regulated PPIs

We found that the extensive interactions of the three antioxidant enzymes with a large number of nuclear proteins, especially those safeguarding the genomic integrity, were up-regulated upon H_2_O_2_ treatment. For example, the interactions of SOD1 with PRIM2, CAT with PRIM2, POLD1 or MCM2 were respectively up-regulated to 130%, 175%, 190%, 144% relative to those treated without H_2_O_2_, as indicated by Western blotting (Fig. 4A and B). In addition, the interactions between SOD1 and CAT or GAPDH, CAT and PRDX2 were respectively up-regulated to 170%, 180% and 139% of control following treatment with H_2_O_2_ (Fig. 4A and B). The integrated analysis of the up-regulated PPIs indicated that a vast majority of the up-regulated PPIs are shared by the three antioxidant enzymes.

**Fig. 4.**
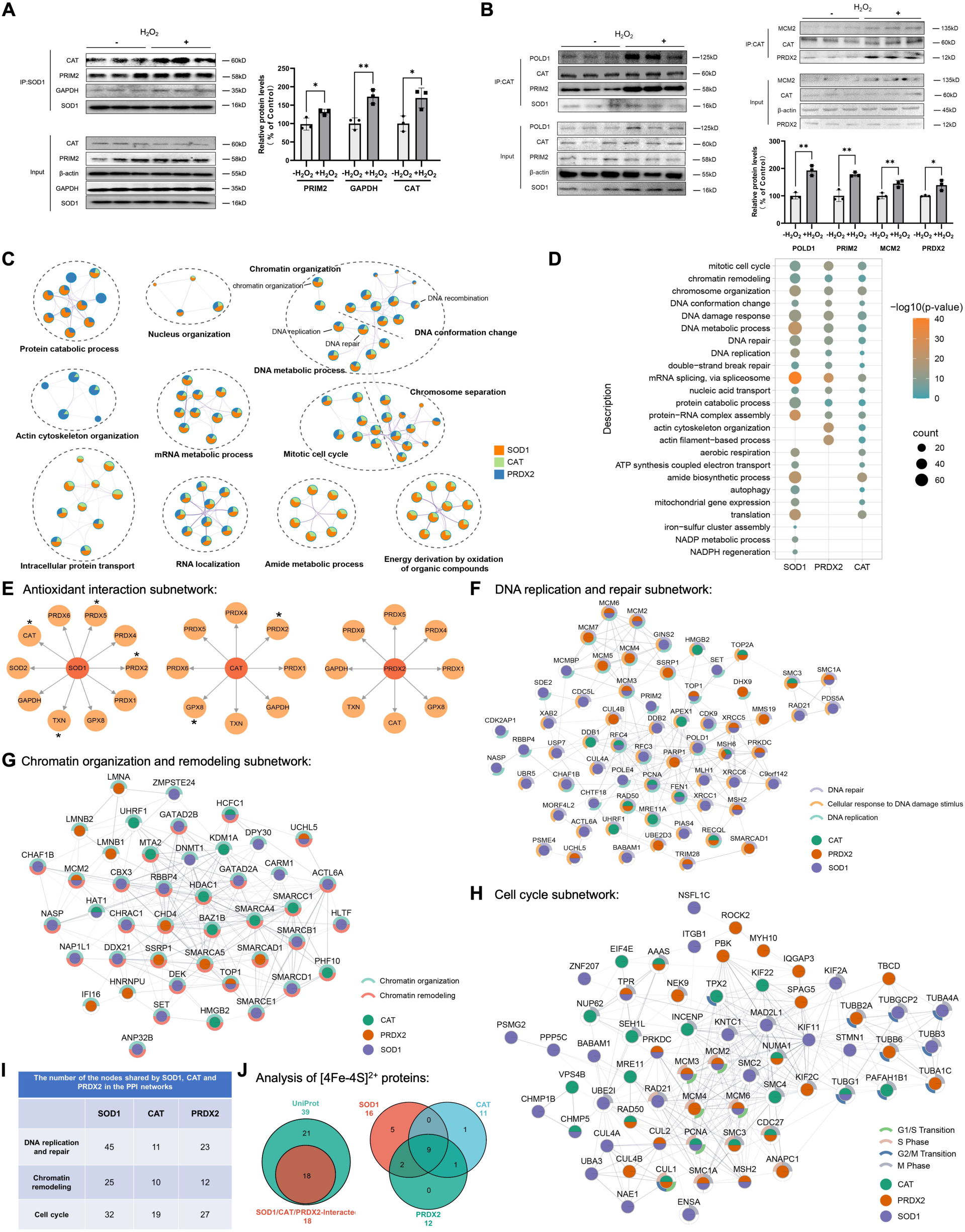
H_2_O_2_ enhances protein subnetworks shared by antioxidant enzymes. **A**) CoIP validation of interactions between SOD1 and CAT/PRIM2/GAPDH. **B**) CoIP validation of interactions between CAT and POLD1/PRIM2/MCM2/PRDX2. **C**, **D**) Functional annotation (**C**) and biological process (**D**) of the proteins whose PPIs with SOD1, CAT or PRDX2 are evaluated by H_2_O_2_ treatment. Protein classification is primarily based on GO annotation. **E**) Antioxidant interaction subnetworks showing the elevated PPIs of antioxidant enzymes by H_2_O_2_ treatment. Asterisks indicate statistical significance of the up-regulated PPIs. **F**-**H**) DNA replication and repair subnetwork (**F**), chromatin organization and remodeling subnetwork (**G**), and cell cycle subnetwork (**H**) showing the elevated PPIs with SOD1, CAT or PRDX2 by H_2_O_2_ treatment. Biological functions are annotated using arc-shaped markers. Subnetworks were constructed by mapping interactomics data onto STRING database. **I**) Number of the nodes shared by SOD1, CAT and PRDX2 in PPI networks. **J**) Number of [4Fe-4S]^2+^ proteins binding to SOD1, CAT and PRDX2. Additionally, an overlap analysis was performed with the [4Fe-4S]^2+^ proteins from UniProt data. For pull-down assays (**A**, **B**), HeLa cells were treated with or without 400 μM H_2_O_2_ for 4 h, and data are means of triplicate samples ± SD (\**P* < 0.05, \*\**P* < 0.01, \*\*\**P* < 0.001; unpaired Student’s t-test). All error bars are SD.

First, the cell functions and processes participated by the up-regulated PPIs were annotated. GO annotation indicated that the PPIs are necessary for the functions including chromatin organization, DNA conformation change, chromosome segregation, mitotic cell cycle, and mRNA metabolism and RNA localization (Fig. 4C). Moreover, the PPIs are requisite for the processes such as DNA metabolism, DNA damage response, DNA replication and repair, chromatin remodeling, chromosome organization, mitotic cell cycle and mRNA splicing (Fig. 4D). Evidently, these functions and processes that occur in the nucleus are essential for maintenance of genomic integrity.

Then, the antioxidant enzymes were observed to form an antioxidant enzyme subnetwork (Fig. 4E), a part of the big network participated by the antioxidant enzymes. Most of the PPIs in the antioxidant enzyme subnetwork were up-regulated upon H_2_O_2_ stimulation. The three antioxidant enzymes were found to interact simultaneously with the other antioxidant enzymes including PRDX1/4/5/6, GPX8, and TXN. PRDXs and GPX8 are essential for reduction of cellular H_2_O_2_ because they promote decomposition of H_2_O_2_, and TXN rescues the inactivated PRDXs.

To understand the role of the up-regulated PPIs in maintenance of genomic integrity, three PPI subnetworks were produced by mapping the interactions between the antioxidant enzymes and the nuclear proteins. In these subnetworks, a large number of overlapped PPIs were found between each antioxidant enzyme and the proteins involved in DNA replication and damage response (Fig. 4F), chromatin remodeling (Fig. 4G) or cell cycle (Fig. 4H), demonstrating the tight association of these PPI subnetworks with the antioxidant enzyme subnetworks (Fig. 4E) and that the enzyme subnetwork is an indispensible module in the antioxidant subnetworks. Further inspection shows that the numbers of the nodes occupied by SOD1 are much more than those by the other two enzymes in the PPI subnetworks, and CAT occupies the minimum network nodes (Fig. 4I), indicating that SOD1 might play a hub role in the formation of these PPI subnetworks. Moreover, the overwhelming majority of the PPIs with the antioxidant enzymes in the PPI subnetworks were up-regulated by H_2_O_2_ treatment, e.g., those of the maintenance proteins of genomic integrity with the antioxidant enzymes, those of MCM components (MCM2/3/6) with SOD1 and PRDX2, those of PCNA or FEN1 with SOD1 and CAT, those of MSH6 with SOD1, CAT and PRDX2. Thus, the PPI subnetworks (Fig. 4, F to H) are indispensible to maintenance of genomic integrity in DNA replication and repair, chromatin remodeling, and cell cycle.

Finally, the interactions between the antioxidant enzymes and the 39 [4Fe-4S]^2+^ proteins found so far in human cells (from UniProt data, 73) were examined. 18 Proteins of them were identified to interact respectively with SOD1 (16), CAT (11) and PRDX2 (12), and 9 proteins were found to interact simultaneously with these three enzymes (Fig. 4J). It is noteworthy that [4Fe-4S]^2+^- containing DNA primase (PRIM2) and polymerase (POLD1) were interact not only with SOD1, but also with CAT and PRDX2, and these PPIs were up-regulated by H_2_O_2_ treatment (Fig. 4F). These indicate that the [4Fe-4S]^2+^ proteins interacted with the antioxidant enzymes play a crucial role in the cellular processes including aerobic respiration, DNA replication and damage response, and iron-sulfur cluster assembly.

The above integrated analysis of three antioxidant interactomes reveals that a big network in the cell is assembled by the three antioxidant enzymes and their interacted partners. Four antioxidant subnetworks, i.e., those are formed by the interactions among antioxidant enzymes, and between the three antioxidant enzymes and the enzymes of DNA replication and damage response, the chromatin remodeling proteins or the cell cycle proteins, are mapped. The vast majority of the PPIs in the four antioxidant subnetworks are significantly up-regulated upon H_2_O_2_ treatment. Therefore, the four subnetworks are indispensible for maintenance of nuclear genomic integrity during oxidative stress.

### The antioxidant networks for maintenance of genomic integrity exist in the nucleus

To further prove the existence of the above-mentioned antioxidant subnetworks for maintenance of genomic integrity (Fig. 4, F to H), an affinity purification-coupled DIA-qMS workflow was designed for examination of the alterations in the interactions between SOD1 and the proteins in the isolated nucleus upon H_2_O_2_ stimulation (Fig. 5A). The nuclear SOD1 interactome was identified with SOD1-HA overexpressed HeLa cells. The PPIs with SOD1 were fixed by addition of formaldehyde (FA) to capture the weak PPIs in the nucleus.

**Fig. 5.**
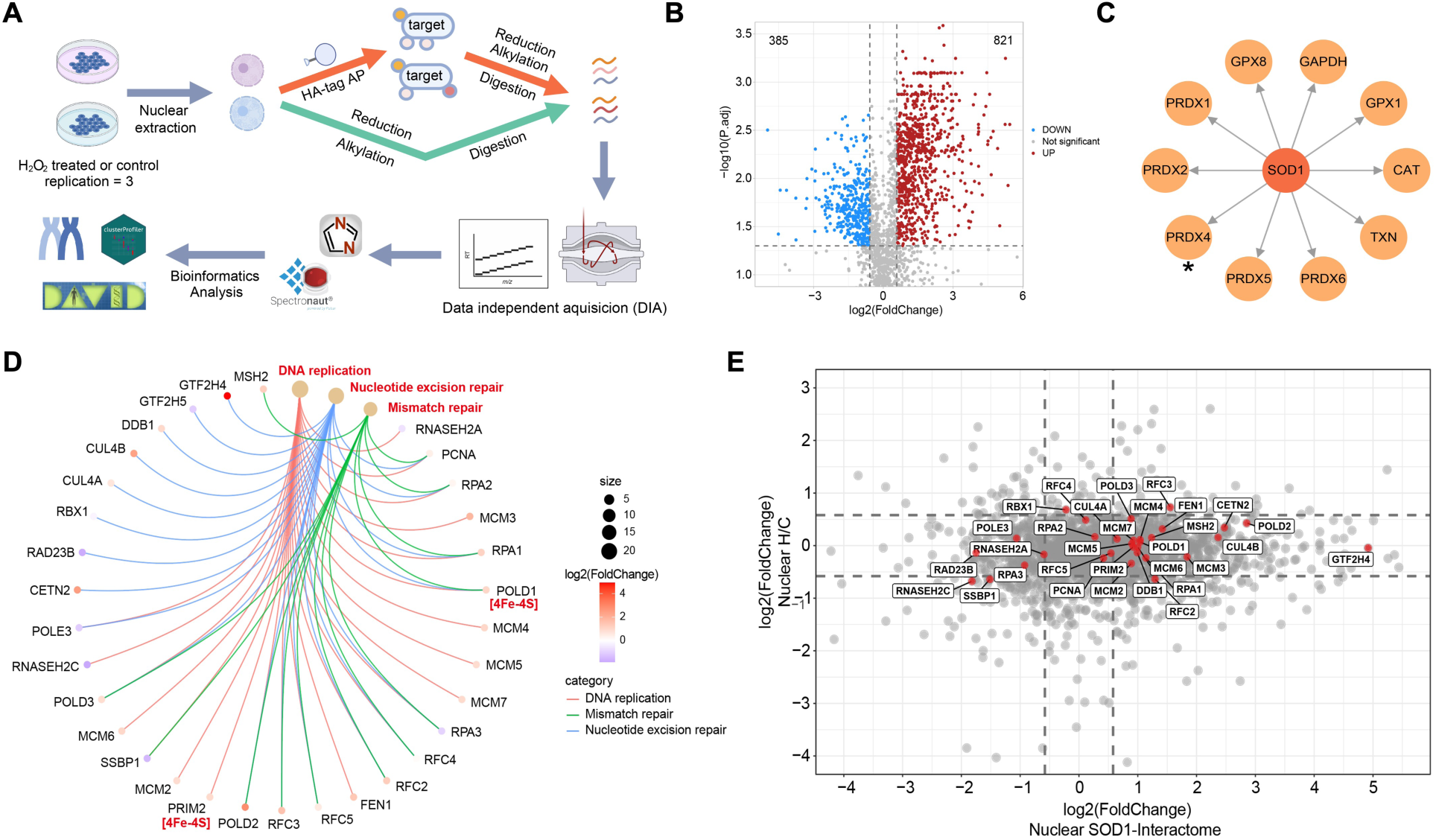
Antioxidant networks that maintain genomic integrity is located in the nucleus. **A**) Flow diagram of nuclear SOD1 interactome by CoIP coupled DIA-qMS. HA-tagged SOD1 (HA-SOD1) was overexpressed in HeLa cells to enhance nuclear SOD1 abundance, followed by quantitative analysis of nuclear proteins using DIA-qMS. **B**) Volcano plot showing the proteins in the nucleus whose interaction with SOD1 are significantly altered by H_2_O_2_ treatment (p. adjust ≤ 0.05, fold change ≥ 1.5). **C**) Antioxidant interaction subnetwork showing the elevated PPIs of antioxidant enzymes with SOD1 by H_2_O_2_ treatment. Asterisk indicates statistical significance. **D**) PPIs between SOD1 and proteins involved in DNA replication, nucleotide excision repair and mismatch repair in the nucleus. **E**) Integrative analysis of the protein quantification (y-axis) and the SOD1 interactome data (x-axis) in the nucleus.

A total of 2268 nuclear proteins were identified to have interactions with SOD1 in the H_2_O_2_- treated and -untreated cells (Fig. 5A). Of these proteins, the interactions of 821 proteins with SOD1 were found to be up-regulated, and those of 385 proteins with SOD1 down-regulated (p. adjust ≤ 0.05, fold change ≤ 1.5) following H_2_O_2_ stimulation (Fig. 5B). Although the number of SOD1-interacting proteins in the nucleus is less than that in the whole cell, the nuclear proteins whose PPIs with SOD1 were altered by H_2_O_2_ stimulation are much more than the proteins in the whole cell, indicating that the SOD1 overexpression allows us to identify much more PPIs with SOD1 than the undisturbed SOD1 expression.

As observed in the whole cell (Fig. 4E), SOD1 interacts not only with CAT and PRDX2, but also with the other antioxidant enzymes with exception of GPX1 in the nucleus (Fig. 5C). The interactions of SOD1 with CAT, PRDXs, GAPDH and TXN, but not GPX8, were up-regulated by H_2_O_2_ treatment. These reveal that the antioxidant enzyme subnetwork (Fig. 4E) is mainly consisted of the PPIs among the antioxidant enzymes in the nucleus.

KEGG enrichment indicated that the elevated PPIs with SOD1 involve in the nuclear pathways including DNA replication and repair, and nucleocytoplasmic transport. The proteins whose PPIs with SOD1 were elevated (fold change >1) are mainly the enzymes essential for DNA replication and repair pathways, e.g., PCNA, FEN, RPA complexes, RFC complexes, MCM complexes, DNA primases and polymerases, MSH and SSBP (Fig. 5D), in agreement with those found in the cell proteome-wide identification of elevated PPIs with SOD1 (Fig. 3G). The [4Fe-4S]^2+^-containing PRIM2 and POLD1 were also found to have the elevated PPIs with SOD1 upon H_2_O_2_ stimulation. These results disclosed that SOD1 involves assembly of an antioxidant subnetwork for maintenance of genomic integrity through the interactions with the nuclear proteins and provide a line of evidence for the existence of the antioxidant subnetwork shown in Fig. 4, E to H.

To probe the factors that lead to the up-regulation of the PPIs with SOD1, the abundance of nuclear proteins in the SOD1-HA overexpressed cell was determined by DIA-qMS (Fig. 5A). The integrated analysis of this determination, together with the nuclear SOD1 interactome data, indicated that 95% of the nuclear proteins whose interactions with SOD1 were altered upon H_2_O_2_ treatment did not display a significant differential in abundance (Fig. 5E, fold change ≤ 1.5). In particular, expression of the proteins in DNA replication and repair was not impacted by H_2_O_2_ treatment. These results are well consistent not only with the cell-wide protein abundance quantification (Fig. 3H), but also with the conclusion that is stemmed from cell-whole proteomic tests of the samples under the conditions of undisturbed and reduced SOD1 expression (Fig. 2). Therefore, the up-regulation of the PPIs with SOD1 can be attributed to the H_2_O_2_-mediated elevation in SOD1 content in the nucleus, which is the same as the conclusion obtained by the determination of the cell-wide SOD1 interactome (Fig. 3H), and also is in line with the recently published global organelle profiling (71).

### The pairwise PPIs in the antioxidant networks were confirmed in the nuclei of single cells

To confirm the existence of the antioxidant subnetworks for maintenance of genomic integrity, quantitative analysis of the pairwise PPIs between the antioxidant enzymes (including GAPDH), and between the antioxidant enzymes and a few of DNA metabolic enzymes were performed *in situ* with proximity ligation assays (PLA, 74). First, the commercial rabbit and goat antibodies against the antioxidant enzymes were observed to emit red immunofluorescence in HeLa cells (fig. S4), indicating that these antibodies can be used in cell immunostaining. Then, the control tests in the absence of primary antibodies, or plus and minus PLA probes for secondary antibodies indicated that the PLA tests are specific to the interactions between the antioxidant enzymes (fig. S9A). Moreover, the PLA signals were not observed, when the cells were treated only with the antibody against the enzymes of interest in the antioxidant enzyme subnetwork (fig. S9B).

The changes in the PLA fluorescence intensity per cell are positively correlated with elevation or reduction in the interactions between two enzymes of interest. The quantitative PLA signals showed that each pair of the enzymes tested interacts to a certain extent whether in H_2_O_2_-untreated cells or in H_2_O_2_-treated cells, but the interactions are significantly up-regulated upon H_2_O_2_ treatment, as illustrated by the elevated PLA signals (Fig. 6A, fig. S10), in alignment with those obtained by Western blotting tests and CoIP-DIA-qMS (Fig. 4, A, B and E). The PLA fluorescence intensity of most of the PPIs in H_2_O_2_-treated cells was more than two-fold of that without H_2_O_2_. These results indicated that the interactions between the enzymes of interest occur in H_2_O_2_- untreated cells, and most of the interactions are up-regulated upon H_2_O_2_ treatment.

**Fig. 6.**
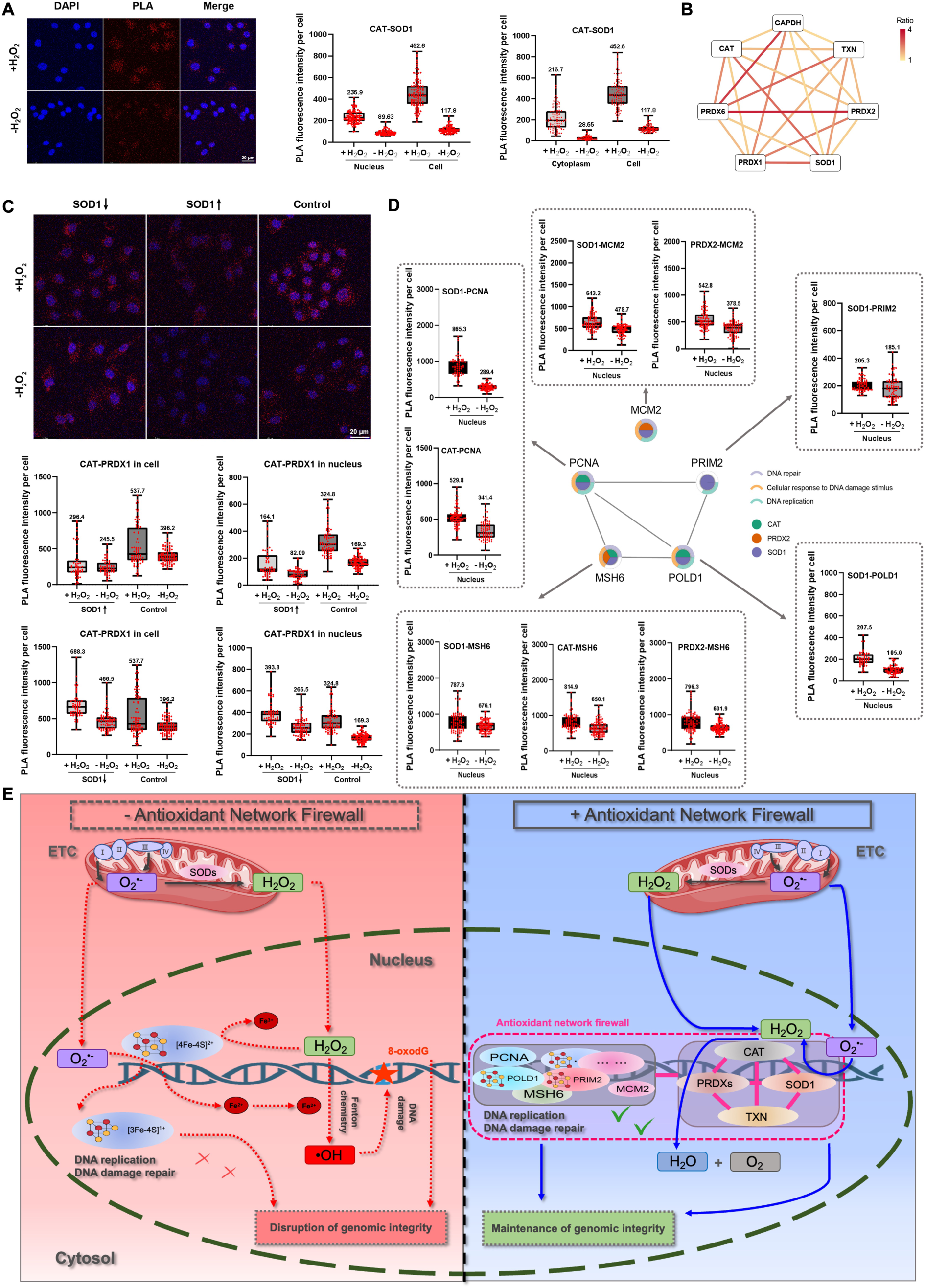
ROS-responsive antioxidant networks, consisting of antioxidant enzymes and nuclear proteins, are present in the nucleus. **A**) PLA visualizing interactions of CAT-SOD1. **B**) Relative PLA signal intensities of pairwise antioxidant enzymes in the nucleus (H_2_O_2_ treatment vs Control). **C**) PLA assay in HeLa cells visualizing SOD1-regulated interactions between CAT and PRDX1. SOD1 expression was modulated *via* overexpression or knockdown. **D**) PLA visualizing interactions between SOD1/CAT/PRDX2 and the proteins involved in maintenance of genomic integrity in the nucleus. For PLA assays (**A**, **C**), HeLa cells were treated with or without 400 μM H_2_O_2_ for 4 h, and stained by a nucleus-specific blue dye DAPI. PLA was performed using specific antioxidant enzyme antibodies. Bar graph depicting the PLA fluorescence intensity per cell (n≥ 50). Representative PLA images from five independent experiments are shown. All error bars are SD. **E**) Proposed mechanistic model for nuclear antioxidant chain firewall. ***Left***: In the absence of a nuclear antioxidant chain, O_2_^•-^ and H_2_O_2_ originated from mitochondria and a nucleus react with [4Fe-4S]^2+^ in DNA-processing proteins. The reactions would destruct [4Fe-4S]^2+^ and produce •OH through Fenton reaction, resulting in not only dysfunction of DNA replication, repair and transcription systems, but also oxidative DNA damage. ***Right***: The antioxidant chain composed of SOD1, CAT, PRDXs and TXN eliminates ROS through catalyzing the decomposition cascade of ROS before they react with [4Fe-4S]^2+^, thereby preventing DNA and DNA-processing proteins from oxidative damage and dysfunction. Thus, the antioxidant chain acts as a firewall in the maintenance of genomic integrity.

We estimated respectively the nuclear and cytoplasmic PLA signal intensity. The PLA signal intensity in the nucleus is stronger than that in the cytoplasm for a majority of the enzyme pairs tested (Fig. 6A, fig. S10). In the H_2_O_2_-treated cells, the PLA signals were also elevated more notably in the nucleus than in the cytoplasm. Indeed, more than half of the nuclear PLA signal intensity was elevated about two-fold upon H_2_O_2_ treatment (Fig. 6B, fig. S10), in line with the Western blotting tests (Fig. 4, A and B). Consequently, although most pairwise PPIs between the antioxidant enzymes were stronger in the nucleus than in the cytoplasm of H_2_O_2_-untreated cells, H_2_O_2_ stimulation significantly enhanced these PPIs both in the nucleus and cytoplasm, as indicated by the described above results (Fig. 4, A, B and E).

The impact of up- and down-regulated SOD1 expression on the pairwise interactions between the antioxidant enzymes (CAT-TXN or -PRDX1, PRDX1-PRDX2 or -PRDX6, TXN-GAPDH) was also examined. In the SOD1-overexpressed cells, these five PPIs were found to be significantly reduced compared to the control, regardless in the whole cell or in the nucleus, whether the cells were treated with H_2_O_2_ or not (Fig. 6C, fig. S11 and S12), as observed in the undisturbed SOD1 expression (Fig. 5A, fig. S10). However, in SOD1 knockdown cells, these PPIs were found to rise both in the whole cell and in the nucleus regardless of H_2_O_2_ treatment, which is in sharp contrast to the SOD1 expression-upregulated and -undisturbed cases. These results indicated that upon SOD1 expression perturbation, the pairwise PPIs between the antioxidant enzymes were altered with nuclear SOD1 levels, and suggested the competitive binding to certain of DNA metabolic enzymes or likely to DNA might occur between SOD1 and other antioxidant enzymes.

On the other hand, we selected the enzymes PRIM2, POLD1 PCNA, MSH6 and MCM2 in the antioxidant subnetworks (Fig. 4, F to H), which are essential for DNA replication and repair, chromatin organization and remodeling, and observed the nuclear PLA signals of interactions between each of them and any of the three antioxidant enzymes. The treatment with H_2_O_2_ results in the up-regulated interactions between SOD1 and each of the selected DNA metabolic enzymes, between CAT and PCNA or MSH6, and between PRDX2 and MSH6, but the down-regulated interactions between CAT or PRDX2 and PRIM2 or POLD1 whether in the whole cell or in the nucleus (Fig. 6D, fig. S13). These altered PPIs were in agreement with those obtained by CoIP-DIA-qMS in the nucleus (Fig. 5, C and D). Obviously, these results further demonstrated that the competitive binding to the DNA metabolic enzymes occurs between the antioxidant enzymes, because the nuclear abundance of the enzymes essential for maintenance of genomic integrity was not significantly altered following H_2_O_2_ treatment (Fig. 5E).

Shortly, PPI mapping performed *in situ* with PLA revealed the elevated PPIs between the antioxidant enzymes, the elevated and reduced PPIs between the antioxidant enzymes and certain of DNA metabolic enzymes upon H_2_O_2_ stimulation. The competitive binding of the antioxidant enzymes to the DNA metabolic enzymes might occur in the nucleus. This additional evidence further supports the presence of the antioxidant subnetworks for maintenance of genomic integrity in the nucleus (Fig. 4, F to H).

## Discussion

In summary, the presence of H_2_O_2_ resulted in oxidative DNA cleavage in the PRIM2-containing solution. The antioxidant enzymes SOD1, CAT and PRDX2 reduced oxidative DNA cleavage in solution by preventing the reaction of H_2_O_2_ with [4Fe-4S]^2+^ in the primase subunit and by the association with DNA. ODD in the human cell exposed to H_2_O_2_ was suppressed by elevating activity of the antioxidant enzymes, but enhanced by reducing their activity. In fact, the whole-cell proteomic determination indicated that reducing SOD1 activity inside the cell compromises the functions of the nuclear proteins essential for maintenance of genomic integrity. These results reveal that the antioxidant enzymes are a guardian of genomic integrity. Integrated interactome analysis of the three antioxidant enzymes indicated the presence of a big antioxidant enzyme-involved PPI network comprising nuclear and cytoplasmic functional modules. Four antioxidant subnetworks, those are formed respectively by the interactions among the antioxidant enzymes and between the antioxidant enzymes and the enzymes responsible for DNA replication and damage response, the chromatin remodeling proteins and the cell cycle proteins, were mapped in the big network. The PPIs among antioxidant enzymes are shared by the four subnetworks. The vast majority of the PPIs in the four subnetworks are significantly up-regulated because of the H_2_O_2_ treatment-mediated nuclear enrichment of the antioxidant enzymes. Therefore, the four antioxidant subnetworks are indispensible for maintenance of genomic integrity during oxidative stress. The identification of the nuclear SOD1 interactome performed with the SOD1 overexpressed cell, together with *in situ* PLA mapping and Western blotting assays of the antioxidant enzyme-involved PPIs in single cells, confirmed the presence of the antioxidant subnetworks in the nucleus. Our results reveal that the antioxidant enzymes safeguard genomic integrity by up-regulating its PPIs with the maintenance proteins of DNA, not by changing abundance of these proteins in the nucleus.

Numerous exogenous stimuli to a cell (75) and nuclear-self endogenous biochemical processes (3–7) can result in increased production of ROS in the nucleus. For example, the treatment of cancer cells with multiple anticancer drugs including platinum-based agents increases nuclear levels of H_2_O_2_ (1), thereby leading to the elevated presence of antioxidant enzymes with direct physical access to DNA in the nucleus (76). H_2_O_2_ is a small electro-neutral molecule that easily reaches any intracellular sites including the interior of a protein molecule. The Fe-S cluster [4Fe-4S]^2+^ was identified as an essential metallocofactor in numerous proteins (69,77–80), but is most vulnerable to oxidative attack from trace intracellular ROS (28,52,54,81). In particular, [4Fe-4S]^2+^ plays a critical role in DNA binding and stabilization of the maintenance proteins and their complexes of genomic integrity (28,67,68,82–87). The cluster in most enzymes of DNA replication and repair (88–90) is located at the sites close to DNA or is exposed to solvents (91). The thermodynamic computation showed that the biding of H_2_O_2_ at [4Fe-4S]^2+^ before reaction is exothermic (92). The reaction of H_2_O_2_ with [4Fe-4S]^2+^ can result in two serious consequences: destruction of the [4Fe-4S]^2+^ in the enzymes and production of •OH through Fenton reaction in the proximity of DNA (Fig. 6E, left), and never stop in the nucleus unless the cellular production of ROS is terminated. The former should be responsible for dysfunction of DNA metabolic enzymes including replication, repair and transcription of DNA, telomere maintenance and chromosome segregation. The most reactive •OH can oxidize the DNA stretches near to the site of its production at a diffusion-controlled rate. In addition, ROS reduces DNA replication fork velocity by causing dissociation of the replisome and even leads to collapse of DNA replication fork integrity (29,30). Moreover, the H_2_O_2_-mediated ODD was observed to be not susceptible to damage repair (93). Therefore, the consequences from the reaction of ROS with [4Fe-4S]^2+^ in the nucleus are loss of both genomic integrity and DNA metabolic functions. The prevention of H_2_O_2_ from destruction of [4Fe-4S]^2+^ in DNA metabolic enzymes and from ODD by •OH is essential for maintenance of genomic integrity. The antioxidant enzymes respectively assemble the antioxidant subnetworks through the PPIs with each other and with the maintenance proteins of genomic integrity, and can block the reaction between ROS and [4Fe-4S]^2+^ through catalyzing the decomposition cascade of O_2_^•-^ and H_2_O_2_ in the nucleus (Fig. 6E, right). The PPIs in the antioxidant subnetworks are up-regulated under oxidative stress and counteract the influence stemmed from oxidative stress on the structures and functions of the DNA maintenance proteins.

Similar to other dynamic nuclear processes, oxidative genome damage was found to change temporally and spatially in the nucleus (13,94), and to associate with gene activation (11), because changes in the intracellular ROS level are time-dependent and regioselective. Moreover, regardless of whether the DNA repair machines are dysfunctional, DNA lesions are prolonged persistent in somatic cells (95,96). Meanwhile, chromatin accessibility and genome organization are also highly dynamic (97,98). The mapping of three antioxidant interactomes indicates the presence of antioxidant subnetworks in the nucleus that are respectively responsible for DNA replication and repair, chromatin remodeling and cell cycle. The PPIs in the subnetworks are up-regulated upon occurrence of oxidative stress. These results are consistent with the dynamic natures of maintenance of genomic integrity. On the other hand, the direct access or binding to DNA and its maintenance proteins of the antioxidant enzymes may be transient and not enter a permanent state. This dynamic antioxidant nature not only facilitates self-regulation of the antioxidant subnetworks to respond to changed ROS levels and to distinct phases of cell cycle, but also enables functional plasticity of the antioxidant enzymes and rapid PPI remodeling in the antioxidant subnetworks under different conditions.

A cell must constantly repair its damaged DNA to accurately and effectively replicate its integral genetic information, because there cannot be any damage to its DNA to be allowed to progress into different phases of cell cycle. The antioxidant subnetworks in the nucleus act as a firewall in maintenance of genomic integrity and might be an alternative mechanism for parent cells to faithfully transfer integral genetic information to offspring cells, in addition to DNA damage response. The antioxidant enzymes in the antioxidant subnetworks can prevent the ROS-mediated destruction of [4Fe-4S]^2+^ by catalyzing the decomposition cascade of O_2_^•-^ and H_2_O_2_ in the nucleus (Fig. 6E, right). This maintenance mechanism of genomic integrity locates at upstream of DNA damage response systems and firstly ensures the intact structures and functions of the [4Fe-4S]^2+^- containing DNA metabolic enzymes. Then, ODD is directly blocked, because the •OH-generating Fenton reaction seldom occurs, thereby reducing the burden of DNA repair machines. Furthermore, since the machines of DNA replication, repair and transcription function smoothly, other types of DNA damage can be effectively repaired in time. It is also possible that the antioxidant subnetworks repress the dioxygen-mediated destabilization of certain Fe-S cluster-containing proteins and their complexes (53), because they can shield the Fe-S clusters from the direct access of small molecules including dioxygen. In addition, the dysfunction of the [4Fe-4S]^2+^-containing DNA replication enzymes caused by regulating PPIs in the antioxidant subnetworks might become a choice for inhibition of viral replication (91). The main limitation of our work is that all tests were performed only using a type of human cancer cells.

## Acknowledgments

We thank Prof. Cuihong Wan and her group (School of Life Science, Central China Normal University) for their help in MS measurements and data analysis.

## Funding

This work was supported by grants from NSFC (21271079, 21771073, 22077046) and self-determined research fund of CCNU from the colleges’ basis research and operation of MOE.

## Author contributions

Conceptualization: C.L. (Changlin Liu); Visualization: H.G., X.L., X.H., Y.C.; Data curation: C.L. (Changlin Liu), H.G., X.H., Y.C., X.L.; Funding acquisition: C.L. (Changlin Liu), X.L.; Writing-original draft: C.L. (Changlin Liu); Writing-review and editing: C.L. (Changlin Liu), H.G., X.L., C.L. (Chunrong Liu); DNA damage tests: X.H., H.C., L.Y., Y.Z. (Yunfeng Zhang); Proteomic identification: H.G., X.L., Y.C., F.W., Y.Z. (Zhao Yao), Y.Y.; PLA: Y.W. All authors read and approved the final manuscript.

## Competing interests

Authors declare that they have no competing interests.

## Data and materials availability

All raw files relating to protein mass-spectrometry are deposited at proteome Xchange, PRIDE number: xxxxxxxx. All other data are available in the main text or the supplementary materials.

## Supplementary Materials

Materials and Methods

Figs. S1 to S13

## Notes

### Competing Interest Statement

The authors have declared no competing interest.

