## Supplementary Materials for "The antioxidant networks act as a firewall in maintenance of genomic integrity"

**The PDF file includes:**

Materials and Methods
Figs. S1 to S13

### Materials and Methods

#### ***1. Cell culture***

HeLa and A549 cells were purchased from China Center for Type Culture Collection (CCTCC) and cultured in DMEM (Gibco, C11995500BT) supplemented with 10% fetal bovine serum (Gibco, A3160802) and 1% penicillin-streptomycin solution (Gibco, 15140122) at 37 $^{o}C$ in a saturated humidity atmosphere containing 5% $CO_2$. HeLa cells stably over-expressing HA-tagged SOD1 were thawed and seeded into 100 mm dishes containing complete medium. Cultures were passaged at least three times to ensure stable growth prior to experimentation. At ~90 % confluence, the medium was removed, monolayers were rinsed once with ice-cold PBS, and 5 mL serum-free DMEM was added. Oxidative stress was induced by adding $H_2O_2$ (10 mM stock) to a final concentration of 0.4 mM. After 4 h at 37 °C, the medium was discarded and cells were fixed with 5 mL PBS containing 1% (v/v) formaldehyde for 10 min at room temperature. Cross-linking was quenched with glycine (0.1 M, 5 min). Cells were washed twice with ice-cold PBS, gently scraped, pelleted at 1 000 × g, flash-frozen in liquid nitrogen, and stored at -80 °C until use.

##### *1.1 Overexpression of SOD1, CAT and PRDX2*

The plasmids encoding human SOD1, CAT, or PRDX2 were cloned into pcDNA3.1(+) vectors. Cells were cultured in 6-well plates for 18-24 h at 70-80% confluent, and transfected with a proper plasmid concentration for 24 h (PRDX2) or 48 h (SOD1 and CAT). Transfection was performed with Lipofectamine 2000 (Invitrogen, 11668019) according to the manufacturer's instructions. HeLa cells were first transfected by plasmids encoding human SOD1 (2.5 μg/mL), CAT (1.25 μg/mL) or PRDX2 (1.25 μg/mL) with increasing content of Lipo2000 (Invitrogen, 11668019) (5-10 μL), and then the protein level was evaluated by western blotting. Optimal transfection conditions were quantitatively analyzed and used in subsequent experiments.

##### *1.2 Knockdown of SOD1, CAT and PRDX2*

siRNAs (siSOD1, siCAT and siPRDX2) were purchased from Genepharma (Shanghai, China), and dissolved in DEPC water at the final concentration of 10 μM. Cells were cultured in 6-well plates for 18-24 h at 40–50% confluent, and transfected with siRNA. Transfection was performed with Lipofectamine RNAiMAX (Invitrogen, 13778075) according to the manufacturer's instructions. HeLa cells were first transfected by siRNA of SOD1 (0-22.5 nM), CAT (0-150 nM) or PRDX2 (0-150 nM), and then the protein level was evaluated by western blotting. Optimal transfection conditions were quantitatively analyzed and used in subsequent experiments. The oligonucleotides were respectively human SOD1 siRNA: 5'-GGCCUGCAUGGAUUCCAUGTT-3', human CAT siRNA: 5'-GGAUCUCACUUGGCGGCAATT-3', human PRDX2 siRNA: 5'-GGAAGTACGTGGTCCTCTT-3'.

#### ***2. Expression, purification and characterization of proteins***

##### *2.1 Expression and purification of antioxidant enzymes*

N-terminally His-tagged human SOD1 and PRDXs were expressed in *E. coli* BL21 (DE3) cells at 37 $^{o}C$ until reaching an $OD_{600}$ of 0.6. Expression of SOD1 and PRDXs was induced by addition of 1 mM IPTG, and cells were cultured at 20 $^{o}C$ for another 20 h. For the induction of SOD1, 0.6 mM $CuSO_4$ and $ZnSO_4$ were co-incubated with the cells. Upon centrifugation, cell pellets derived from 1 L bacterial culture were resuspended in 30 mL lysis buffer (20 mM Tris-HCl, 150 mM NaCl, 10 mM PMSF, pH 8.0-8.5), sonicated, and centrifuged. The supernatant was applied to a Nickel-NTA charomatography column, washed with 2-3 column volumes of buffer A (50 mM Tris-HCl, 10 mM imidazole, pH 8.0~9.0) and buffer B (50 mM Tris-HCl, 30 mM imidazole, pH 8.0-9.0), and bound proteins were eluted with the elution buffer (50 mM Tris-HCl 250 mM imidazole, pH 8.0-9.0) in 2-3 column volumes. Further purification was performed using SP-Sepharose (GE Healthcare) columns.

N-terminally $His_6$-SUMO-tagged human TXN was expressed in *E. Coli* BL DE3 cells at 37 $^oC$ until reaching an $OD_{600}$ of 0.6. Expression of TXN was induced by addition of 0.5 mM IPTG, and cells were cultured at 20 $^oC$ for another 12 h. Upon centrifugation, cell pellets derived from 1 L bacterial culture were resuspended in 30 mL lysis buffer (20 mM HEPES, 150 mM NaCl, 10 mM PMSF, pH 7.0), sonicated, and centrifuged. The supernatant was applied to a Nickel-NTA charomatography column, washed with 2-3 column volumes of buffer C (20 mM HEPES, 10 mM imidazole, pH 7.0) and buffer D (20 mM HEPES, 30 mM imidazole, pH 7.0), and bound proteins were eluted with the elution buffer (20 mM HEPES 250 mM imidazole, pH 7.0) in 2-3 column volumes. Finally, 20 U/mg of thrombin was used to cleave the label of $His_6$-SUMO, and the proteins were purified again by Nickel-NTA charomatography column. Further purification was performed using SP-Sepharose (GE Healthcare) columns.

#### *2.2 SOD1 activity assay*

SOD1 activity was determined by SOD Assay Kit (Sigma, 19160-1KT-F) according to the manufacture's protocol. Briefly, cells were seeded in 6-well plates for 18-24 h, subjected to indicated treatment, and lysed by lysis buffer (Beyotime, P0013) containing PMSF. 80 μL of ice-cold chloroform/ethanol (37.5/62.5 (v/v)) was added into 50 μL lysate, shaked for 30 s and centrifuged at 2500 ×g for 10 min. 20 μL lysate with equal protein content was mixed with a 200 μL WST working solution and a 20 μL enzyme working solution, and incubated for 20 min at 37 $^oC$. The absorbance of resulted water-soluble WST-1 formazan at 450 nm was collected using SpectraMax M5. Relative SOD1 activity was calculated according to the manufacturer's instructions.

#### *2.3 Catalase activity assay*

Catalase activity was determined by catalase Assay Kit (Beyotime, S0051) according to the manufacture's protocol. Briefly, cells were seeded in 6-well plates for 18-24 h, subjected to indicated treatment, and lysed by cell extraction buffer containing PMSF. 3 μL lysate with 6 μg protein was first mixed with 37 μL catalase working solution, and incubated with 10 μL 250 mM $H_2O_2$ at 25 $^oC$ for 3 min, then incubated with 450 μL reaction-stop solution. 10 μL above-mentioned solution was first mixed with 40 μL catalase working solution, and then 10 μL mixture was incubated with 200 μL fresh color-substrate solution for 18 min at 25 $^oC$. The absorbance at 450 nm was collected using SpectraMax M5. Relative catalase activity was calculated according to the protocol.

#### *2.4 Expression and purification of PRIM2*

The PRIM2 plasmid was transformed into *E. coli* TOP10 Super Competent Cells, and the bacterial culture was inoculated onto solid medium supplemented with ampicillin and chloramphenicol, followed by incubation at 37 °C. Single colonies were selected for activation and expanded until reaching an $OD_{600}$ of 0.6. Protein expression was induced by adding 0.5 mM

IPTG, 100 μg/mL ferric ammonium citrate and 5 μM $ZnSO_4$, and the culture was incubated at 16 °C with shaking at 180 rpm for 12-16 h. The bacterial cells were harvested by centrifugation, and the pellet was resuspended in lysis buffer (50 mM phosphate, 1 M NaCl, 5% glycerol, 5 mM imidazole, 1 mM $MgCl_2$, pH 8.0). Next, PMSF (300 μM), DNase (1 μg/mL), and lysozyme (4 μg/mL) were added to the suspension for lysis. After centrifugation, the supernatant was loaded onto a pre-equilibrated Ni-NTA column and washed sequentially with the following buffers: 50 mM phosphate, 1 M NaCl, 5% glycerol, 10 mM imidazole, 1 mM $MgCl_2$, pH 8.0; 50 mM HEPES, 0.3 M NaCl, 5% glycerol, 20 mM imidazole, 1 mM $MgCl_2$, pH 8.0; 50 mM HEPES, 0.3 M NaCl, 5% glycerol, 250 mM imidazole, 1 mM $MgCl_2$, pH 8.0.

Under an argon atmosphere in glovebox, 17 μM of the purified PRIM2 without $[4Fe\text{-}4S]^{2+}$ was treated with 850 μM DTT for 5 min to establish a reducing environment. Ferrous ammonium sulfate (FAS, 34 μM) and $Na_2S$ (34 μM) were added stepwise: FAS was added and incubated for 2 min; $Na_2S$ was added and incubated for 5 min. Next, the reaction was incubated on ice in the dark for 5 h, and then EDTA (68 μM) was incubated with the mixture for 5 min under argon to chelate residual FAS and centrifugation to remove precipitates. Finally, PRIM2 with $[4Fe\text{-}4S]^{2+}$ was desalted using a desalting column.

*2.5 Characterization of PRIM2 by inductively coupled plasma massspectrometry*

Copper and sulfur contents of PRIM2 were detected by inductively coupled plasma massspectrometry (ICP-MS). In short, digestion was performed by mixing 400 μL protein stock solution with 500 μL $HNO_3$, followed by microwave-assisted sonication at 60 °C for 2 h. After filtration through a 0.45 μm membrane, the sample was diluted to 10 mL and subjected to ICP-MS analysis. Instrument parameters (Thermo, iCAP™ RQ ICP-MS) were set as follows: emission power at 1150 W, argon pressure at 0.6 MPa, gas flow rate at 0.7 L/min, and detector as CID.

### 3. Agarose gel electrophoresis

Agarose gel (1%) was prepared by dissolving agarose in 1 × TAE buffer (pH~8), cooling to 40°C and adding 4S Green Nucleic Acid Stain. After solidification, the gel was submerged in 1 × TAE buffer. Next, PRIM2, plasmid DNA, $H_2O_2$ and antioxidant enzymes were mixed and incubated at 37 °C for 30 min. After diluted with 6 × DNA loading buffer. Electrophoresis was performed at 160 V for 20 min in 1 × TAE buffer. Images were captured using a gel imaging system.

### 4. DNA damage testing in vivo

*4.1 Cell viability assay*

About 2000 cells per well containing 200 μL DMEM were plated on a 96-well, cultured overnight, and subjected to indicated treatments. 20 μL Cell Counting Kit-8 (CCK-8) (Biosharp, BS350) was added to each well, and the cells were further incubated for another 1 h. The corrected absorbance of 450 nm was measured by SpectraMax M5 and normalized to the control group.

*4.2 Comet assay*

Cells were seeded in 6-well plates for 18-24 h, subjected to indicated treatment, digested and resuspended with 1 mL ice-cold PBS. 30-50 μL cell suspension was mixed with 700 μL low-melting agarose gel, spotted onto frosted glass slides coated with agarose gel, and solidified at

4 °C. Slides were immersed into pre-chilled lysis buffer (0.5 M EDTA, 12.5 μg/mL Proteinase K, 2% N-Lauroylsarcosine sodium salt) at 4 °C in the dark for 4 h, to lyse the cells embedded in agarose gel. After cell lysis, slides were used for electrophoresis in 1 × TBE buffer with a voltage at 1 volt/cm for 30 min at room temperature, and incubated with 40 μg/mL propidium iodide for 0.5-1 h in the dark. The images were visualized under Olympus microscope. Statistical analysis was performed by CometScore 2.0.

*4.3 Quantitative ELISA assay of 8-OHdG*

Human 8-Hydroxy-desoxyguanosine (8-OHdG) from HeLa cells was quantified by ELISA according to the manufacturer's instructions (Bioswamp, HM10783). Briefly, cells were seeded in 6-well plates for 18-24 h and subjected to indicated treatment. Total DNA was extracted using commercial kit (Omega, D3396-02). 40 μL solution with equal content of DNA was incubated with 10 μL biotinylated 8-OHdG antibody and 50 μL HRP-Conjugate Reagent at 37 ºC for 30 min. After washing three times, Chromogen Solution A (50 μL) and B (50 μL) were incubated with DNA at 37 ºC for 10 min, and then mixed with 50 μL Stop Solution. The 450 nm absorbance was collected using SpectraMax M5. Relative 8-OHdG level was calculated according to the manufacturer's instructions.

***5. Measurement of intracellular $H_2O_2$ levels***

To determine the level of intracellular $H_2O_2$, cells were incubated with the final concentration of 10 μM 2′,7′-dichlorofluorescin diacetate (Sigma, D6883) for 20 min at 37°C in the dark, washed three times with PBS, and resuspended in 200 μL PBS. Flow cytometry (BD, Accuri™ C6) was used to determine the fluorescent intensity.

***6. Proximity ligation assay***

All the buffers and solutions were provided in the Duolink® in-situ PLA kit (Sigma, DUO92008). First, 400 μL of 0.3% Triton X-100 was added to each sample for permeabilization treatment for 10 min, and then the samples were washed twice with PBS for 5 min. Next, the cells were blocked with 40 μL Duolink® blocking solution for 1 h at 37 °C, in a humidity-controlled chamber. After removing the blocking solution, antibodies were incubated with the cells in a 37 °C incubator for 1 h. Subsequently, PLUS (8 μL) and MINUS (8 μL) secondary antibody probes were mixed and diluted with 24 μL Duolink® Antibody Diluent. After washing the samples with Wash Buffer A, the cells were incubated with the mixed secondary antibody probes at 37 °C for 1 h, and co-incubated with 1 × ligase solution at 37 °C for 30 min. Under dark conditions, the polymerase was diluted with 1 × Duolink® Amplification Buffer, followed by incubation with cells at 37 °C for 100 min. Finally, the slides were treated with Duolink® In Situ DAPI Mounting Medium for 15 min. All images were acquired on a Leica TCS-SP8. The excitation sources were set as 405 nm (DAPI), 488 nm (Alexa Fluor 488), and 594 nm (Texas Red). Z-stacks were acquired and maximal projection images (2048 × 2048 pixels) were analyze using ImageJ. The fluorescence intensity of PLA signals per cell was provided by multiplying the mean fluorescence intensity (the total cell fluorescence intensity is first subtracted by its background fluorescence intensity, and then divided by the cell area) of the cell by the area of its PLA signals. The PLA signals per cell were imaged at least 5 times under a confocal microscope and processed, and the PLA fluorescence intensity of at least 60 randomly selected cells was respectively determined for the interactions of each enzyme pair and its control.

### *7. Characterization of protein and mRNA levels*

#### *7.1 Western blotting*

Proteins were separated by SDS/PAGE electrophoresis and transferred to polyvinylidene fluoride (PVDF) (Thermo Scientific, 88585) according to proven technique. PVDF membranes were blocked in 1xTBS (Thermo Scientific, 28360) buffer with 5% BSA for 1 h. A suitable concentration of primary antibodies diluted by 1xTBS was incubated with the membranes overnight at 4 °C, followed by 2 h incubation at room temperature with secondary antibodies (Bopsharp, BL003A). The HRP-DAB Substrate Kit (TIANGEN, PA110) was used for chromogenic reaction. Following acquisition, images were imported into Image J for quantification. The primary antibodies used were anti-SOD1 (Abcam, ab252426, 1:1000 dilution), anti-CAT (Abcam, ab76024, 1:1000 dilution), anti-PRDX2 (Abcam, ab109367, 1:1000 dilution), anti-PRDX6 (Abcam, ab59543, 1:1000 dilution), anti-TXN (Abcam, ab26320, 1:1000 dilution), anti-ACTB (Proteintech, 20536-1-AP, 1:1000 dilution), anti-GAPDH (Proteintech, 10494-1-AP, 1:1000 dilution).

#### *7.2 RT-qPCR*

Total RNAs were isolated using the High Pure RNA Isolation Kit (Roche, 11828665001). RNA was reverse-transcribed using the Transcriptor First Strand cDNA Synthesis Kit (Roche, 04897030001). RT-qPCR was performed using the FastStart Essential DNA Green Master (Roche, 06402712001). All experiments were performed according to the manufacturer's instructions. The results were calculated by the $2^{-\Delta\Delta Ct}$ method and matched to control samples. All primers were SOD1 sense primer: 5′-AAGGCCGTGTGCGTGCTGAA-3′, SOD1 antisense primer: 5′-GGCCCACCGTGTTTTCTGGA-3′); CAT sense primer: 5′-CCAGAAGAAAGCGGTCAAGAA-3′, CAT antisense primer: 5′-GAGATCCGGACTGCACAAAG-3′; PRDX2 sense primer: 5′-CACCTGGCTTGGATCAACACC-3′, PRDX2 antisense primer: 5′-CAGCACGCCGTAATCCTCAG-3′; GAPDH sense primer: 5′-CGGAGTCAACGGATTTGGTCGTAT-3′, GAPDH antisense primer: 5′-AGCCTTCTCCATGGTGGTGAAGAC-3′.

### *8. Proteomics testing*

#### *8.1 Nuclear protein extraction*

A frozen pellet (~1 × $10^8$ HA-SOD1–expressing cells) was thawed on ice and resuspended in 500 μL Cytoplasmic Extraction Reagent A supplemented with 1 % (v/v) protease-inhibitor cocktail. The suspension was vortexed for 10 s and rested for 10 s, repeating this cycle for 2 min, followed by 30 min on ice. Lysates were centrifuged (10 000 × g, 5 min, 4 °C) and the supernatant discarded. The pellet was resuspended in 500 μL lysis buffer with 1 % protease inhibitors, sonicated to disrupt nuclei, and centrifuged (12 000 × g, 10 min, 4 °C). The resulting nuclear protein fraction was collected for immediate use or stored at -80 °C.

#### *8.2 Nuclear immunoprecipitation of SOD1*

Anti-HA magnetic beads (40 μL) were equilibrated as above, then incubated overnight at 4 °C with 500 μL nuclear lysate. Beads were washed five times with lysis buffer, and bound proteins were eluted twice with 400 μL elution buffer (10 min each, room temperature). Pooled eluates were dried for downstream analysis. Three biological replicates were processed per condition.

*8.3 Enrichment of proteins binding to antioxidant enzyme*

Protein A/G beads (100 µL) were equilibrated as described. Frozen pellets ($2 \times 10^7$ cells) were lysed in 500 µL lysis buffer with 1 % protease inhibitors and sonication. Antibody (30 µL) was added, and lysates were incubated overnight at 4 °C with rotation. Complexes were captured on pre-washed beads for 2 h at room temperature, washed five times, and eluted twice with 400 µL elution buffer. Combined eluates were dried. Three biological replicates were obtained for each condition.

*8.4 Data-dependent acquisition (DDA) spectral-library preparation*

A single 100 mm dish of frozen cells was lysed in 1 mL lysis buffer containing 1 % protease inhibitors. Lysates were sonicated on ice and cleared (12 000 × g, 5 min, 4 °C). Protein concentration was determined by BCA and adjusted to 1 mg $mL^{-1}$. Aliquots (300 µL) underwent methanol-chloroform precipitation. Pellets were dissolved in 300 µL 8 M urea (ammonium bicarbonate). Disulfide bonds were reduced with TCEP (5 mM, 30 min, 25 °C) and alkylated with IAA (10 mM, 30 min, dark). Samples were diluted with 1.2 mL 50 mM ammonium bicarbonate, and sequencing-grade trypsin was added at 1:50 (w/w). Digestion proceeded overnight at 37 °C.

Digested peptides were desalted on a C18 SPE cartridge, eluted with 80 % ACN/10 % methanol, and dried. Peptides were reconstituted in 100 µL basic-pH solvent A (5 mM ammonium formate, pH 10) and fractionated on a preparative C18 column at 1 mL $min^{-1}$ using the following gradient of solvent B (5 mM ammonium formate in 90 % ACN, pH 10): 0–7 min, 0 % B; 7–13 min, 0–16 % B; 13–73 min, 16–40 % B; 73–77 min, 40–44 % B; 77–82 min, 44–60 % B; 82–96 min, 60–90 % B; 96–105 min, 90 % B. Fractions were collected as specified and dried.

Each fraction was resuspended in 30 µL solvent A (0.1 % formic acid) and 1 µL was injected onto a nano-UHPLC coupled to a high-resolution Orbitrap mass spectrometer. The analytical gradient (solvent B, 0.1 % formic acid in 80 % ACN) was: 0 min, 1 % B; 10 min, 5 % B; 10–70 min, 5–20 % B; 70–90 min, 20–35 % B; 90–97 min, 35–95 % B; 97–101 min, 95 % B; 101–103 min, 50 % B at 500 nL $min^{-1}$; 103–107 min, 50 % B; 107–110 min, 2 % B at 300 nL $min^{-1}$.

For whole-cell SOD1 interactomes, peptides were acquired on an Orbitrap Fusion Lumos with the following settings: spray voltage 2.2 kV; capillary 320 °C. $MS^1$: resolution 120 000 (m/z 200), AGC $2 \times 10^5$, m/z 350–1 800, max injection 110 ms. DDA $MS^2$: resolution 30 000, AGC $5 \times 10^4$, max injection 54 ms, cycle 2 s, isolation window 1.3 m/z, stepped HCD 20/25/30 %, dynamic exclusion 30 s (charge states 1 and >6 excluded, intensity $\geq 4 \times 10^4$).

For nuclear SOD1, CAT, and PRDX2 interactomes, peptides were acquired on an Orbitrap Q-Exactive under analogous parameters ($MS^1$ resolution 70 000; $MS^2$ resolution 17 500; isolation window 1.6 m/z; other settings as above). Raw files from the 12 fractions were imported into Spectronaut 14 (Biognosys). A DDA spectral library was generated with default parameters and subsequently used for DIA analysis.

*8.5 Data-independent acquisition (DIA) and data analysis of SOD1, CAT and PRDX2 interactomes*

Freeze-dried affinity-purified samples were resuspended in 100 µL ice-cold water, mixed with 600 µL pre-chilled (−20 °C) acetone, and precipitated overnight at −20 °C. Pellets were collected (12 000 × g, 5 min, 4 °C), dissolved in 100 µL 8 M urea (ammonium bicarbonate), reduced with TCEP (5 mM, 30 min, 25 °C), alkylated with IAA (10 mM, 30 min, dark), diluted with 400 µL

50 mM ammonium bicarbonate, and digested overnight at 37 °C with trypsin (0.1 mg $mL^{-1}$). Peptides were desalted and dried for LC-MS/MS.

Peptide loading and the nano-LC gradient matched the DDA method. For whole-cell SOD1 interactomes, DIA data were acquired on an Orbitrap Fusion Lumos: $MS^1$ resolution 120 000, AGC $2 \times 10^5$, m/z 350–1 200; $MS^2$ resolution 30 000, AGC $1 \times 10^5$, 50 variable isolation windows spanning m/z 350–1 800. For nuclear SOD1, CAT and PRDX2 interactomes, data were acquired on an Orbitrap Q-Exactive: $MS^1$ resolution 70 000, AGC $3 \times 10^6$, m/z 350–1 800; $MS^2$ resolution 17 500, AGC $1 \times 10^6$, 20 fixed windows covering m/z 400–900.

Spectra were searched in Spectronaut 14 using the "DDA library + Direct" workflow with default settings. Proteins exhibiting differential expression according to Spectronaut (adjusted p-value ⩽ 0.05, fold change ⩾ 1.5) underwent functional annotation and enrichment analysis using the clusterProfiler R package (v4.0) and Metascape.

#### *8.6 DIA and data analysis of proteomes in cells with $H_2O_2$ treatment and SOD1 disturbance*

Protein isolation and tryptic-peptide generation for cells with different $H_2O_2$ treatment and SOD1 expression were performed exactly as described for spectral-library (DDA) construction. Subsequent data-independent acquisition followed the same nano-LC gradient and DIA settings specified for the nuclear SOD1, CAT, and PRDX2 interactome analyses.

Spectra were searched in DIA-NN 1.8.1 using the "DDA library + Direct" workflow with default settings. The protein expression matrix was imported into Perseus software (version 1.9). Proteins exhibiting a missing value rate below 30% across all samples were retained. Missing values for these retained proteins were replaced with values sampled from a normal distribution. Differentially expressed proteins were identified using one-way ANOVA with a False Discovery Rate (FDR) threshold of < 0.05. Proteins showing significant differential expression underwent z-score normalization and were subsequently subjected to hierarchical clustering analysis. Proteins residing within clusters of interest were then functionally annotated using Metascape.

### ***9. Statistical analysis***

Statistical analyses were performed using GraphPad Prism (version 8, GraphPad Software). For pairwise comparison, unpaired Student's *t* test was used to determine statistical significance. For multiple-group comparison, One-way ANOVA test was utilized to determine significance. Error bars in all graphs represent SD from at least three independent experiments. *P* or adjusted *P* values less than 0.05 (**P* < 0.05, ***P* < 0.01, ****P* < 0.001) were considered statistically significant.

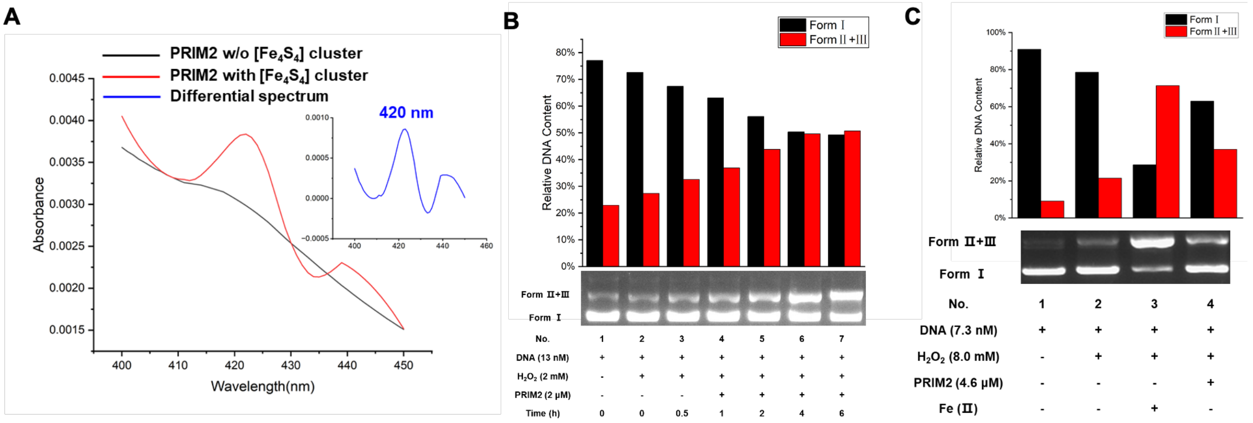


**Fig. S1. Characterization of $[4Fe\text{-}4S]^{2+}$ and oxidative DNA cleavage in solution. A**) UV-vis absorption spectroscopy showing the presence of $[4Fe\text{-}4S]^{2+}$ in reconstituted PRIM2. **B**) Agarose gel electrophoresis showed that $H_2O_2$ mediates time-dependent (0-6 h) cleavage of plasmid DNA, in the presence of PRIM2 protein containing $[4Fe\text{-}4S]^{2+}$. **C**) Agarose gel electrophoresis showed that $H_2O_2$ disrupts $[4Fe\text{-}4S]^{2+}$clusters in PRIM2 and releases iron to enhance oxidative DNA cleavage. Representative electrophoresis images are presented, and quantitative results were derived from grayscale analysis of DNA bands.

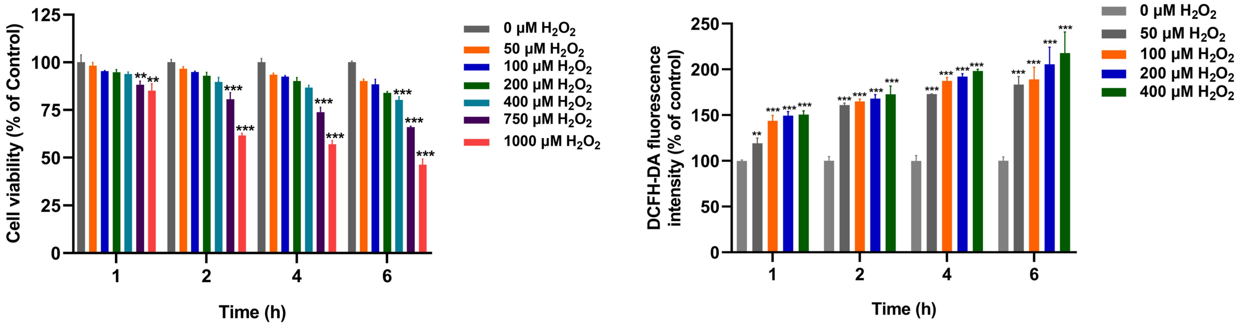


**Fig. S2. $H_2O_2$ uptake and its impacts on cell viability. A**) Dose- and time-dependent effects of $H_2O_2$ (0-1000 μM incubation for 1-6 hours) on the viability of HeLa cells, assessed by MTT assay. **B**) Intracellular $H_2O_2$ levels of HeLa cells. The cells were treated with $H_2O_2$ (0-400 μM) for 1-6 hours. Relative intracellular $H_2O_2$ levels were measured using DCFH-DA, a $H_2O_2$ fluorescent probe. Data are means of triplicate samples ± SD (*$P < 0.05$, **$P < 0.01$, ***$P < 0.001$; One-way ANOVA). All error bars are SD.

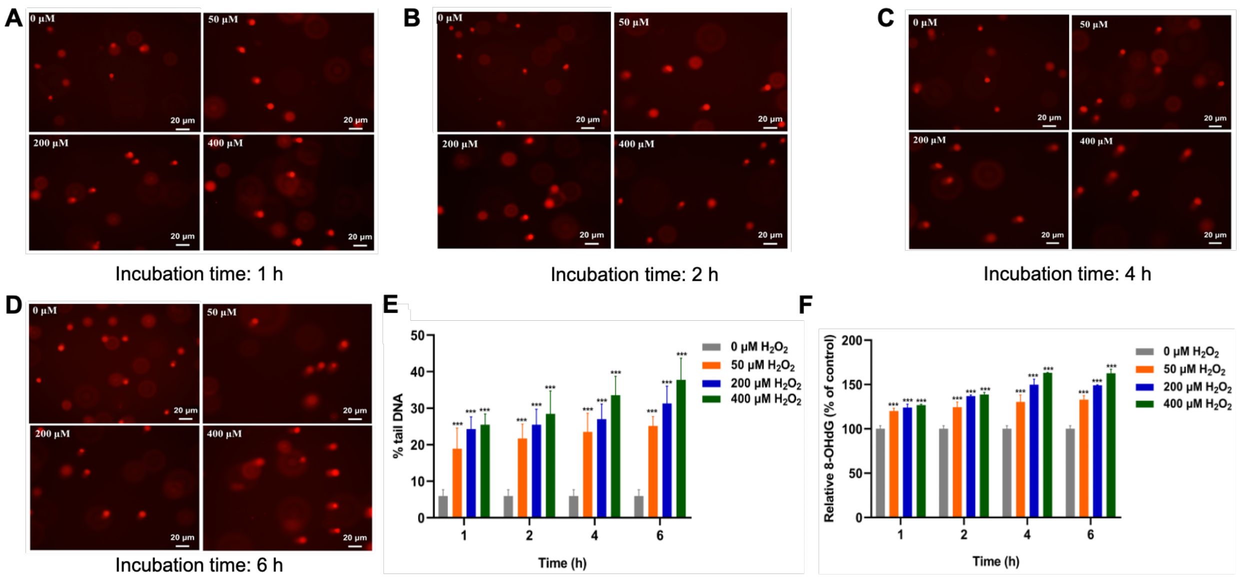


**Fig. S3. $H_2O_2$-induced oxidative DNA damage. A-D**) Representative comet assay images showing oxidative DNA damage by the treatment with $H_2O_2$. **E**) Quantitative analysis of DNA tail content in comet assay. **F**) Detection of 8-OHdG, a marker of oxidative DNA damage. Here, HeLa cells were incubated with 0-400 µM $H_2O_2$ for 1-6 hours. Data are means of triplicate samples ± SD (**P* < 0.05, ***P* < 0.01, ****P* < 0.001; One-way ANOVA). All error bars are SD.

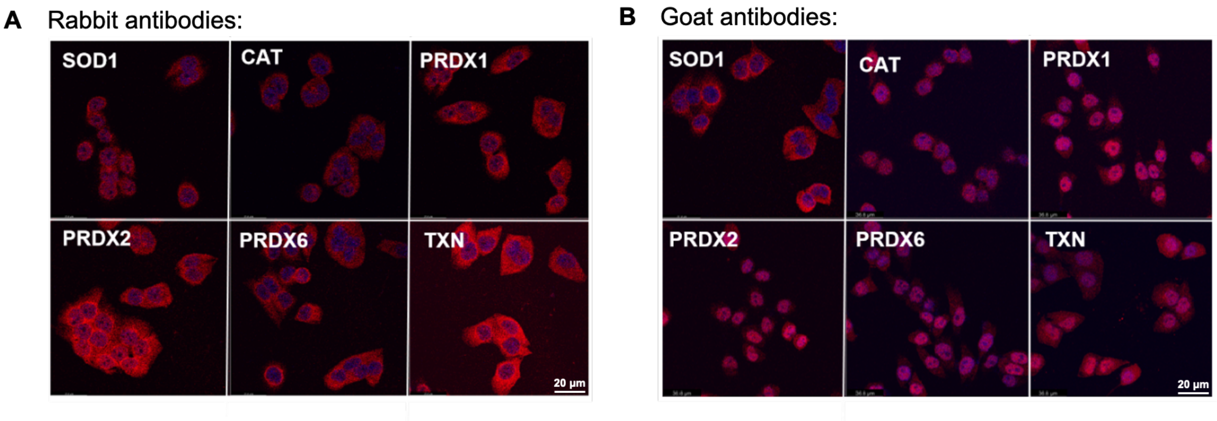


**Fig. S4. Subcellular localization of antioxidant enzymes in HeLa cells.** *In situ* visualization of antioxidant enzymes by immunofluorescence using antibodies derived from distinct species. The nucleus was stained by DAPI, a blue dye for nucleus. Scale bar: 20 µm. Representative results from three independent experiments are shown.

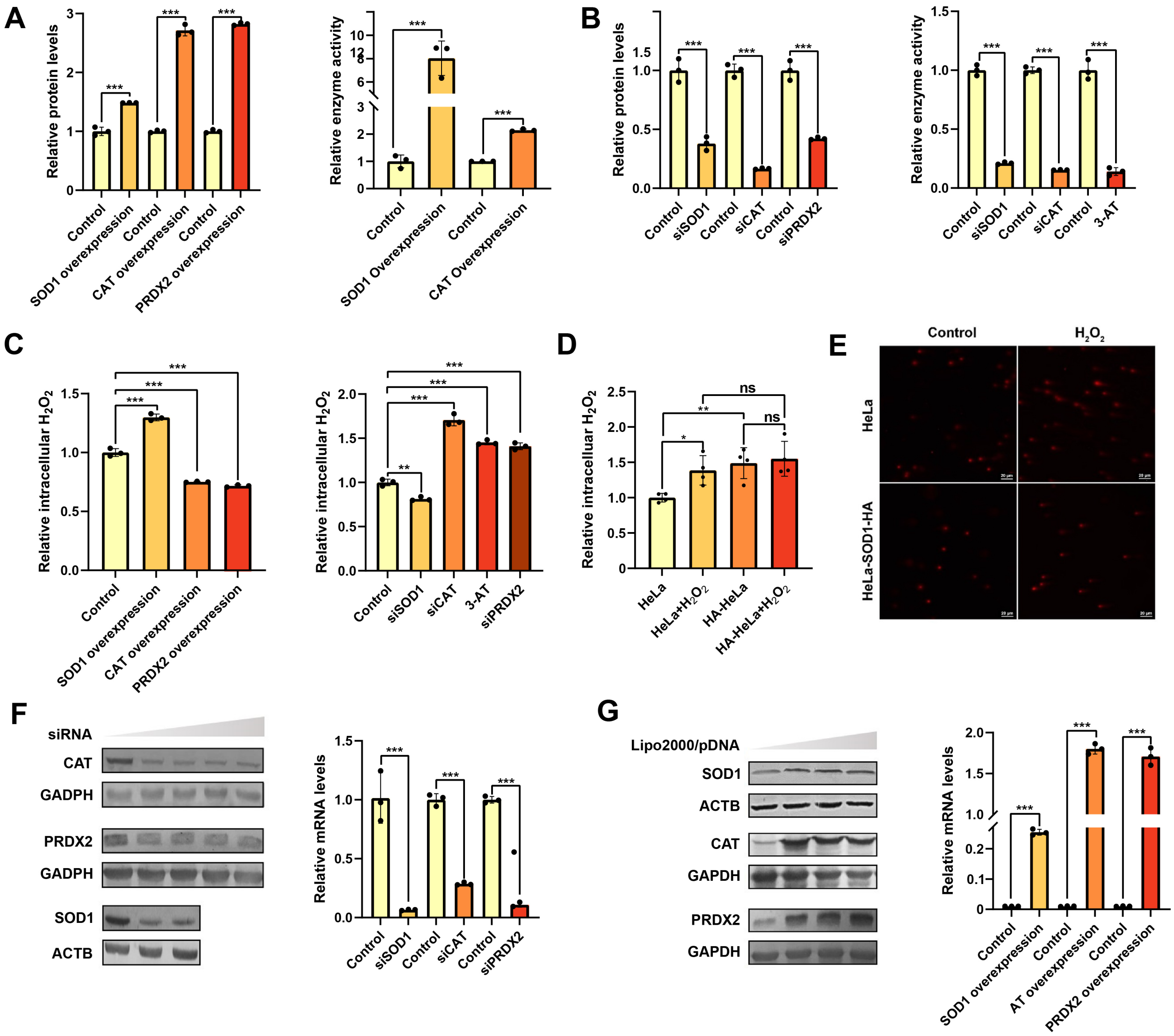


**Fig. S5. Regulation of antioxidant enzyme expression and the impacts on intracellular $H_2O_2$ levels. A, B, F, G**) Following overexpression/knockdown/inhibition of SOD1, CAT and PRDX2, the protein/mRNA/activity levels were assessed by Western blotting, RT-qPCR and commercial kits, respectively. Representative results from three independent experiments are presented. **C, D**) Quantification of intracellular $H_2O_2$ levels. $H_2O_2$ treatment was conducted at a concentration of 400 µM for 4 h. For CAT inhibition, HeLa cells were incubated with 10 mM **3-AT** (1,2,4-Triazole). Data are means of triplicate samples ± SD (*$P$ < 0.05, **$P$ < 0.01, ***$P$ < 0.001; unpaired Student's t-test), and all error bars are SD. **E**) Comet tail assays showed that SOD1 overexpression mitigates oxidative DNA damage. HeLa cells were treated with or without 400 µM $H_2O_2$, and harvested for comet assays.

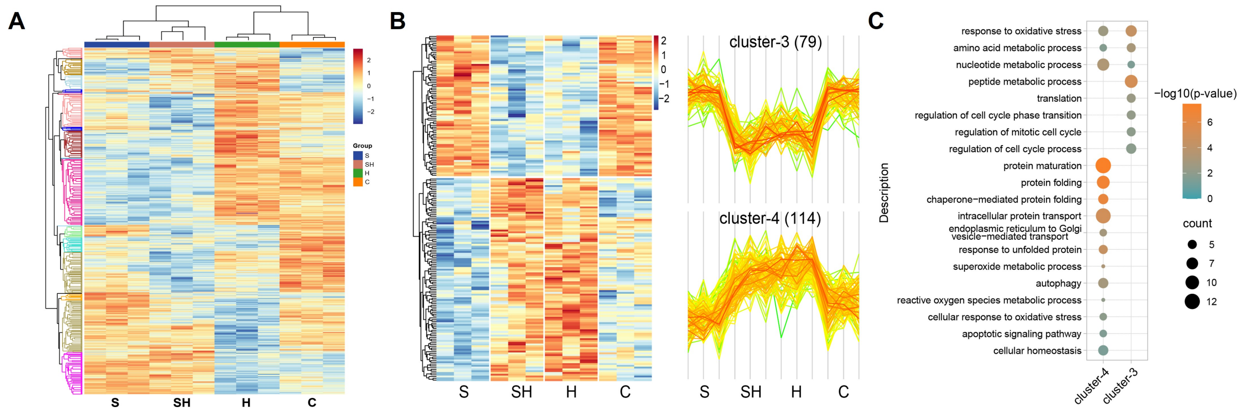


**Fig. S6. Clustering analysis of differentially expressed proteins by SOD1 knockdown and $H_2O_2$ treatment.** **A**) Clustering analysis of all differentially expressed proteins (**C**: control, **H**: $H_2O_2$ treatment, **S**: SOD1 knockdown, **SH**: SOD1 knockdown and $H_2O_2$ treatment). **B**, **C**) Clustering (**B**) and GO analysis (**C**) of proteins specifically responsive to $H_2O_2$ treatment. Protein abundance was normalized based on MS signal intensity (**A**, **B**).

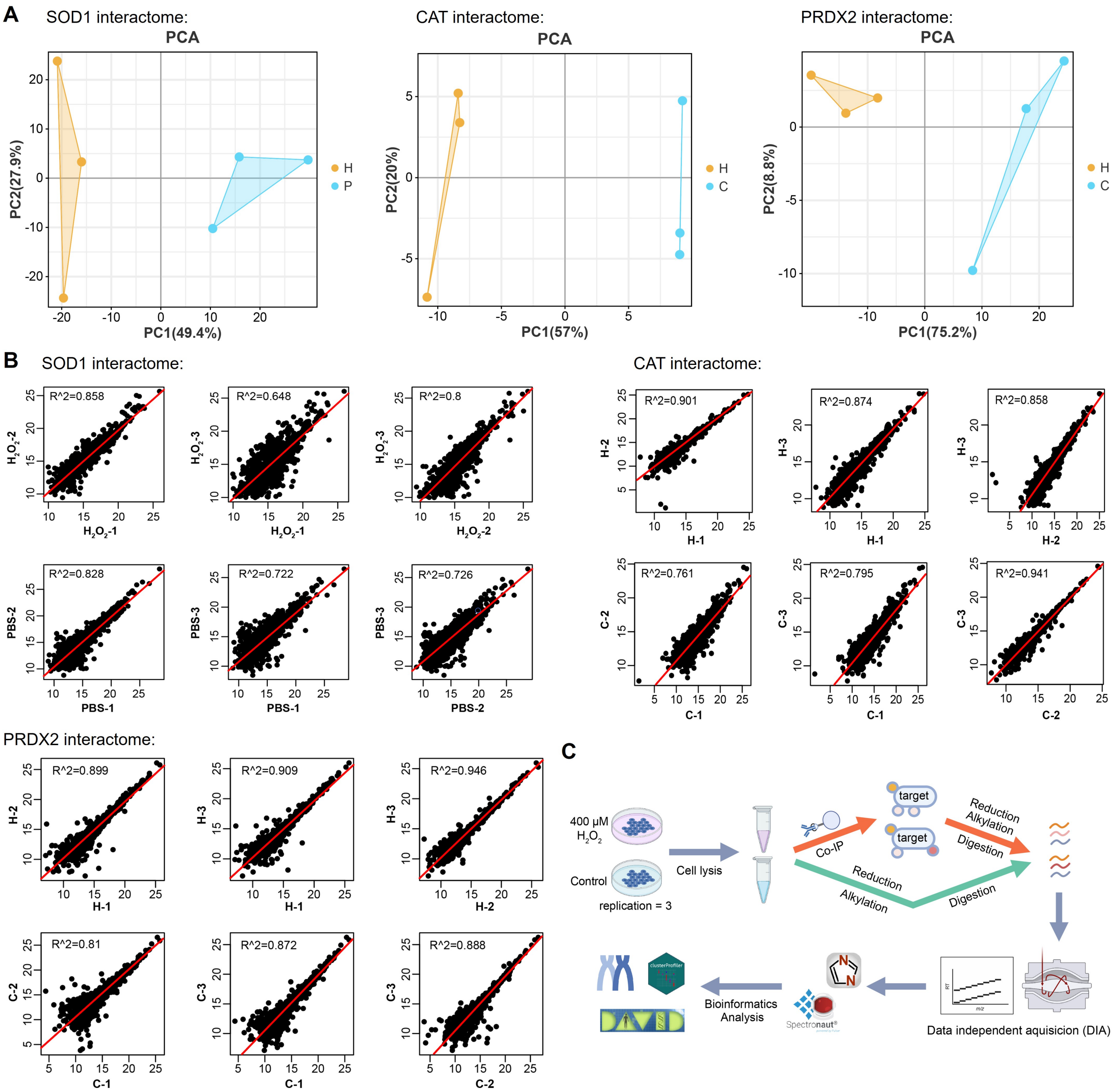


**Fig. S7. Quality assessment of the antioxidant interactomes. A**) PCA analysis of the antioxidant interactomes. **B**) Correlation analysis of replicate samples within the antioxidant interactomes. All quality assessments were performed using quantitative proteomic data form the interactomes. **C**) Flow diagram for the identification of the antioxidant interactomes and for the quantification of whole-cell proteins.

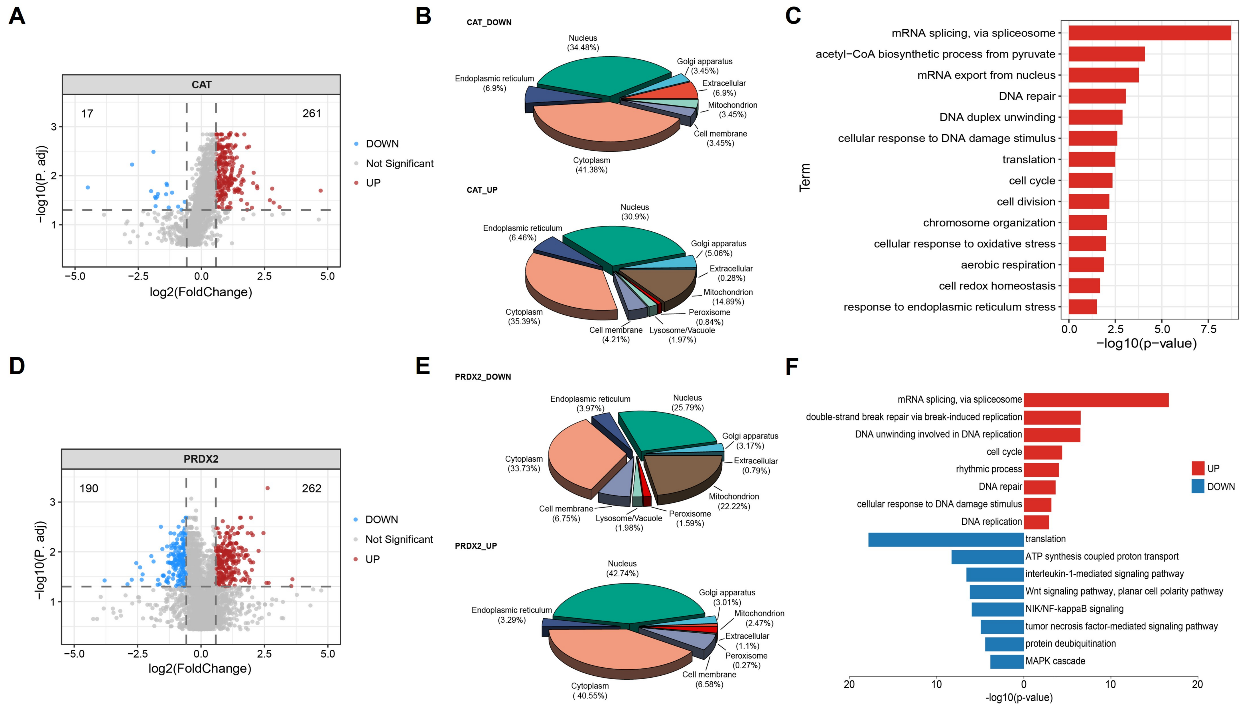


**Fig. S8. Characterization and functional annotation of CAT and PRDX2 interactomes. A**, **D**) Volcano plot depicting the proteins whose interaction with CAT (**A**) and PRDX2 (**D**) are significantly altered upon $H_2O_2$ treatment (p. adjust ≤ 0.05, fold change ≥ 1.5). **B**, **C**) Subcellular localization (**B**) and GO analysis (**C**) of the proteins whose PPIs with CAT were respectively up- and down-regulated by $H_2O_2$ treatment. **E**, **F**) Subcellular localization (**E**) and GO analysis (**F**) of the proteins whose PPIs with PRDX2 were respectively up- and down-regulated by $H_2O_2$ treatment.

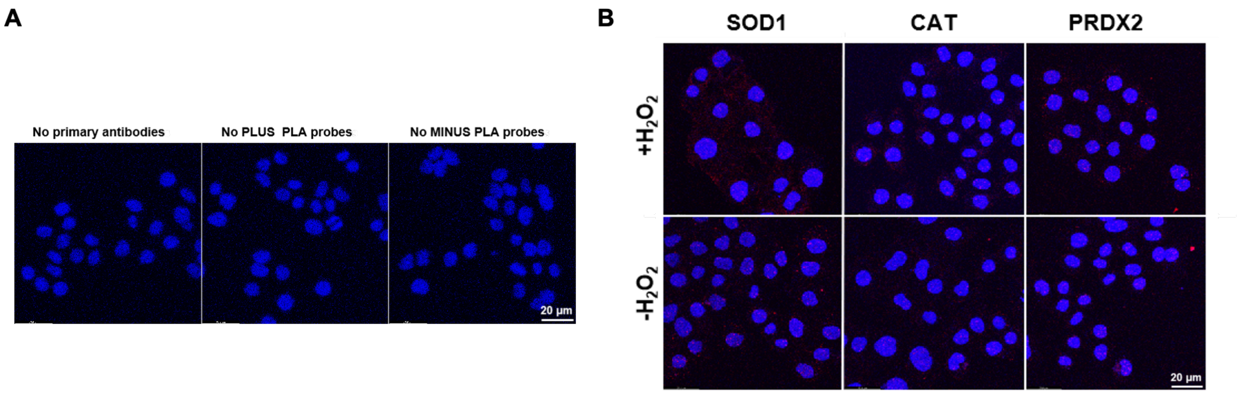


**Fig. S9. Feasibility assessment of *in situ* PLA for detecting protein-protein interactions. A**) Evaluation of PLA background signals in the absence of primary antibodies, PLA probes and MINUS PLA probes. **B**) Characterization of PLA background signals with the addition of antioxidant enzyme antibodies. HeLa cells were treated with or without 400 μM $H_2O_2$ for 4 h, and the nucleus was stained by DAPI. Scale bar: 20 μm. Representative results from three independent experiments are shown.

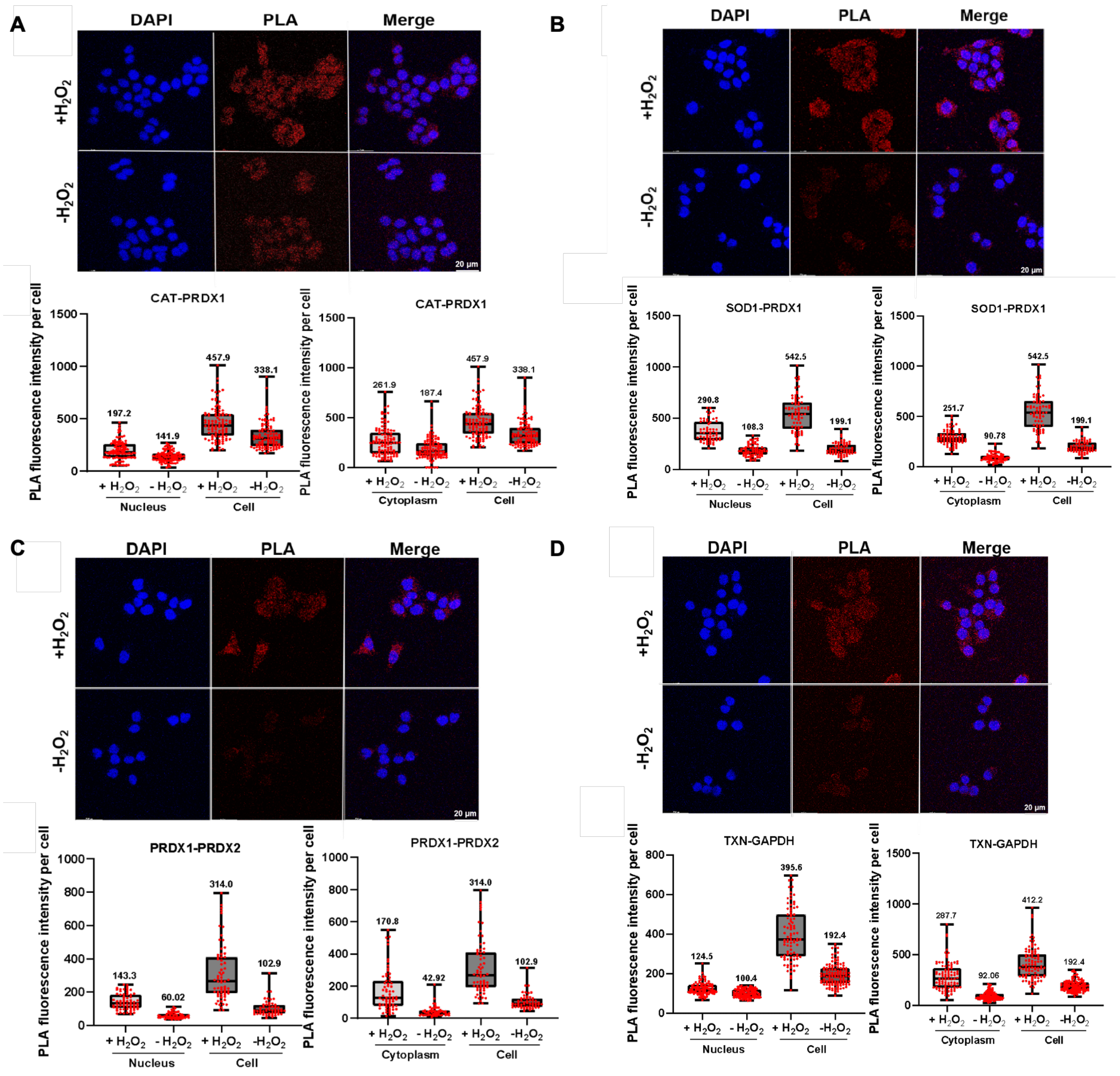


**Fig. S10. Characterization of antioxidant enzyme interactions by *in situ* PLA. A-D**) PLA visualizing interactions of CAT-PRDX1 (**A**), SOD1-PRDX1 (**B**), PRDX1-PRDX2 (**C**) and TXN-GAPDH (**D**). HeLa cells were treated with or without 400 μM $H_2O_2$ for 4 h, and stained by a nucleus-specific blue dye DAPI. PLA was performed using specific antioxidant enzyme antibodies. Bar graph depicting the PLA fluorescence intensity per cell in the nucleus and the cytoplasm (n⩾50). Representative PLA images from five independent experiments are shown. All error bars are SD.

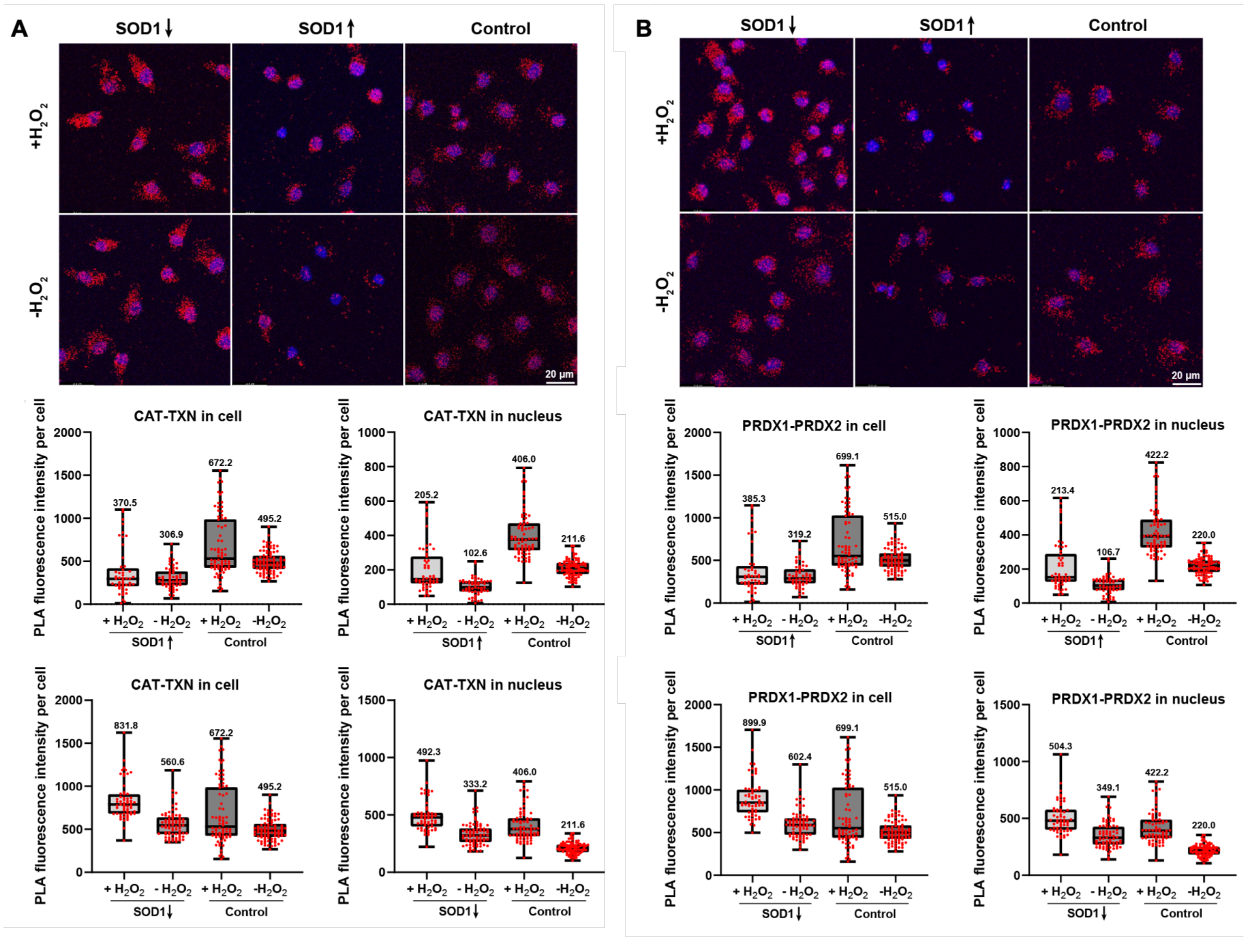


**Fig. S11. SOD1 modulates antioxidant enzyme interactions in the nucleus. A**, **B**) PLA visualizing interactions of CAT-TXN (**A**), and PRDX1-PRDX2 (**B**) regulated by SOD1. HeLa cells were treated with or without 400 μM $H_2O_2$ for 4 h after overexpression or knockdown of SOD1. The cells were then stained by a nucleus-specific blue dye DAPI. PLA was performed using specific antioxidant enzyme antibodies. Bar graph depicting the PLA fluorescence intensity per cell in the nucleus and the whole cell (n⩾50). Representative PLA images from five independent experiments are shown. All error bars are SD.

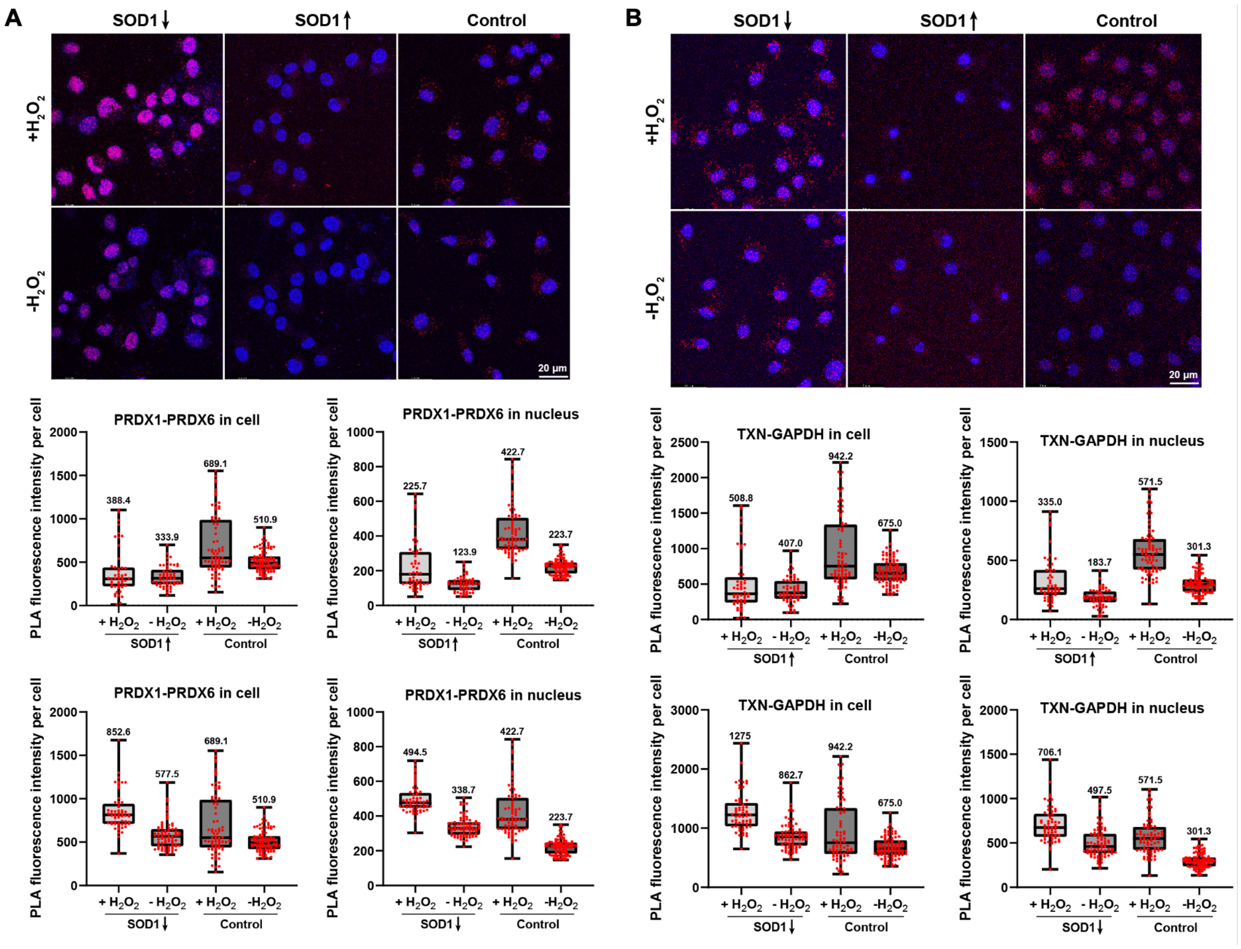


**Fig. S12. SOD1 modulates antioxidant enzyme interactions in the nucleus. A**, **B**) PLA visualizing interactions of PRDX1-PRDX6 (**A**), and TXN-GAPDH (**B**) regulated by SOD1. HeLa cells were treated with or without 400 μM $H_2O_2$ for 4 h after overexpression or knockdown of SOD1. The cells were then stained by a nucleus-specific blue dye DAPI. PLA was performed using specific antioxidant enzyme antibodies. Bar graph depicting the PLA fluorescence intensity per cell in the nucleus and the whole cell (n⩾50). Representative PLA images from five independent experiments are shown. All error bars are SD.

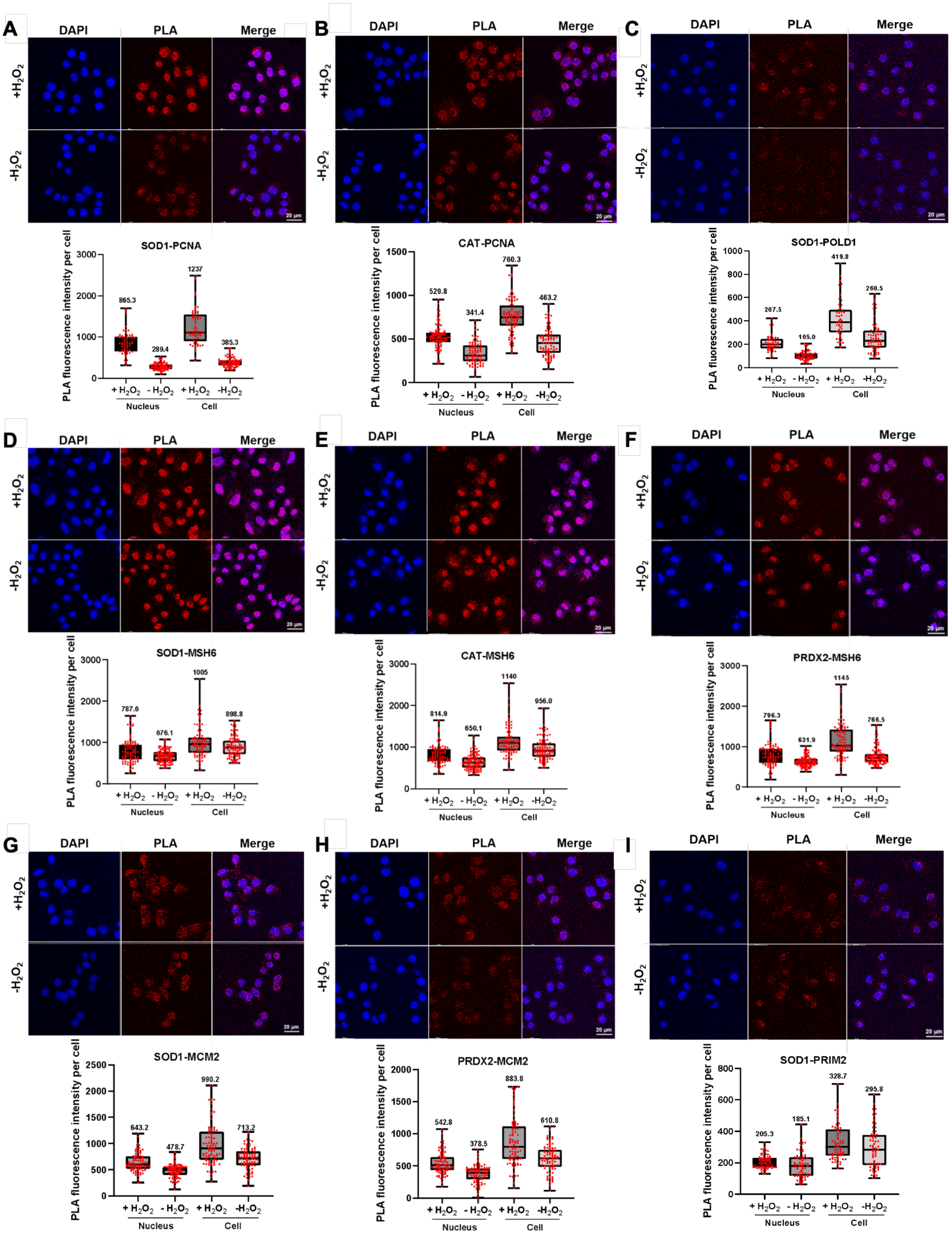


**Fig. S13. The interactions between antioxidant enzymes and nuclear proteins are regulated by $H_2O_2$. A-I**) PLA visualizing interactions of SOD1-PCNA (**A**), CAT-PCNA (**B**), SOD1-POLD1 (**C**), SOD1-MSH6 (**D**), CAT-MSH6 (**E**), PRDX2-MSH6 (**F**), SOD1-MCM2 (**G**),

PRDX2-MCM2 (**H**) and SOD1-PRIM2 (**I**) in the nucleus. HeLa cells were treated with or without 400 μM $H_2O_2$ for 4 h, and stained by a nucleus-specific blue dye DAPI. PLA was performed using specific antibodies. Bar graph depicting the PLA fluorescence intensity per cell in the nucleus and the whole cell (n⩾50). Representative PLA images from five independent experiments are shown. All error bars are SD.
